# Novel *Bacillus altitudinis* endolysin (ArtE2) targeting both Gram-positive and Gram-negative bacteria

**DOI:** 10.64898/2026.01.19.700289

**Authors:** Roxana Portieles, Xinmin Ma, Jianjian Hu, Hongli Xu, Xiangyou Gao, Nayanci Portal González, Rabia Durrani, Ramon Santos-Bermúdez, Orlando Borrás-Hidalgo

## Abstract

Antibiotic resistance is a major global health concern. The development of new antibiotics and therapeutics is crucial for the future. Bacteriophages produce endolysins that induce bacterial lysis, making them a promising treatment option. Deep sequencing was used to identify and isolate genes encoding endolysins from *Bacillus* spp.. We characterized the biological activities of these endolysins. Additionally, this study focused on the design of a new chimeric endolysin, ArtE2, which combines endolysin-2 with a polycationic peptide to address the bacterial activity in gram-negative pathogenic bacteria. Using bioinformatic tools, we conducted three-dimensional modeling of the endolysin ArtE2 and its interactions with peptidoglycan fragments. In this study, we tested the activity of chimeric endolysins against both Gram-positive and Gram-negative bacteria. All endolysins share the same catalytic domain and diverse cell-binding domains. Some endolysins are highly specific to certain bacterial species or strains, whereas others have broader specificities. Histidine interactions are an important part of the mechanism by which ArtE2 connects with bacterial peptidoglycan. Additionally, the engineered endolysin ArtE2 was highly effective at killing *Staphylococcus aureus* and *Escherichia coli*. In silico analysis showed that the fusion did not negatively affect endolysin folding or activity. These findings suggest that ArtE2 could be used to develop efficient antibacterial controls targeting pathogenic Gram-positive and Gram-negative bacteria.

## INTRODUCTION

Antibiotic discovery, development, and use represent important scientific milestones for human survival, saving lives every year since their discovery (1–3). The use of both natural and synthetic versions has proven beneficial for combating extremely virulent and deadly bacteria (4, 5). The overuse of preventative measures rather than therapy and treatment of viral infections, along with the nonspecificity of antibiotics, has resulted in the development of resistant bacterial strains, creating a huge global health concern (6, 7).

The ineffectiveness of antibiotics in treating infections caused by resistant bacteria results in a significant number of deaths annually (8–10). The silent pandemic is predicted to escalate in the coming years owing to the increasing number of deaths caused by drug-resistant bacteria (9). The rise of antibiotic-resistant pathogens such as methicillin-resistant *Staphylococcus aureus* (MRSA), vancomycin-resistant *Enterococcus* (VRE), multidrug-resistant *Mycobacterium tuberculosis* (MDR-TB), third-generation cephalosporin-resistant *Escherichia coli* (3GCREC), third-generation cephalosporin-resistant *Klebsiella pneumoniae* (3GCRKP), carbapenem-resistant *K. pneumoniae* (CRKP), carbapenem-resistant *Pseudomonas aeruginosa* (CRPA), and carbapenem-resistant *Acinetobacter baumannii* (CRAB) constitutes a global challenge in health systems worldwide (11, 12).

Despite the development of new and effective antibiotics, this issue remains unresolved. The World Health Organization has implemented different approaches to combat resistance, usually through new mechanisms of action (12). To manage this problem, rapid diagnosis, knowledge of resistance mechanisms, avoidance of antibiotic use in animal feed, international collaboration, and implementation of solutions through research and development are essential (13). Development of innovative therapeutic strategies for the treatment of bacterial infections is a crucial step in the future.

Bacteriophages are viruses that selectively target and kill bacteria (14). This organism produces a battery of enzymes during its life cycle (15, 16). Among these enzymes, endolysins hydrolyze the host cell wall and subsequently allow for the release of bacteriophage progeny. Consequently, these enzymes are key components of the phage cycle and good alternatives to antibiotics (17–21).

Endolysin activity is classified into different classes, including acetylmuramidases, transglycosylases, glucosaminidases, amidases, and endopeptidases (20). In addition, endolysins have one or two (multi-domain) N-terminal enzymatically active domains (EADs) with a short, flexible linker motif associated with the C-terminal cell wall-binding domain (CBD). This structure is found in gram-positive bacteriophages (22). The EADs of endolysins cleave diverse specific peptidoglycan bonds in the murein layer of host bacteria, whereas CBDs recognize and bind to different components of the cell wall for the proper activity of EADs (22). However, endolysins of phages infecting gram-negative hosts have simple EADs without CBDs and with one or two CBDs at the N-terminus and EADs at the C-terminus (23).

Endolysins, which can lyse gram-positive bacteria, have been studied as potential antimicrobial agents (24, 25). These have shown efficacy in eliminating pathogenic bacteria from mucosal surfaces and treating systemic infections (24). Their effectiveness against antibiotic-resistant bacteria, lack of toxicity, and synergy with antibiotics makes them promising for the treatment of infectious diseases (26). Previous studies have shown that endolysins can be used in food safety, pathogen detection, disinfection, nanotechnology, and vaccine development (25, 27–31).

The antibacterial action of endolysins in eradicating bacterial pathogens was first limited to pathogenic gram-positive bacteria such as *Streptococcus* sp., *Bacillus anthracis*, *Enterococcus* sp., and *Staphylococcus aureus* (24). This benefit is due to the permeability of the peptidoglycan layer in gram-positive bacteria (32). However, the outer membrane of gram-negative bacteria hinders the effectiveness of endolysins, which cannot penetrate easily (21, 33, 34).

Currently, these enzymes have been engineered to overcome this limitation (33–35). The use of permeabilizing peptides promotes the transfer of the fusion enzyme across the outer membrane (33, 36). After translocation, peptidoglycans are degraded, which causes rapid cell death via osmotic lysis. Modification of endolysins using molecular biology tools and computational modeling aims to enhance their antimicrobial capacity, stability, and effective delivery, which are crucial for their clinical applications (35). Although less common than antibiotics, there is a theoretical risk of bacteria developing resistance; however, their specificity often mitigates this risk (37).

Previously, a group of Bacillus endolysins has been identified and characterized (insert reference). Many of these endolysins exhibit relatively broad antimicrobial activities (38, 39). Recently, the endolysin PlyJBA6 from *Bacillus amyloliquefaciens* was shown to lyse gram-negative bacteria without destabilizing their outer membranes (40). Regarding the structure of Bacillus, a study revealed that several endolysin domains remain experimentally uncharacterized. Investigating the features of these domains will help us understand how endolysins can be used to develop new antibacterial drugs (41).

The use of endolysins is a promising strategy for the treatment of bacterial infections. Here, we report the identification, characterization, and biological analysis of *Bacillus altitudinis* and *Bacillus aryabhattai*. In this study, an engineered endolysin (ArtE2) was designed, analyzed, expressed, and characterized against Gram-positive and Gram-negative bacteria. Interestingly, the endolysin ArtE2 efficiently controlled Gram-positive and Gram-negative bacteria in combination with a polycationic peptide.

## MATERIALS AND METHODS

### Bacterial strains and growth conditions

*Escherichia coli* BL21 (DE3) was used for cloning and protein synthesis. Antimicrobial activity assays were performed using *E. coli* BL21 (DE3) and *Staphylococcus aureus* (Bmb939) grown in Luria-Bertani (LB) medium (Thermo Fisher Scientific, US) aerobically at 37 °C with constant shaking at 200 rpm. The stocks of bacterial strains were stored in 20% glycerol at -80 °C.

### Sequence analysis of *B. altitudinis* and *B. aryabhattai* endolysins

Sequences encoding endolysins present in the genomes of *B. altitudinis* and *B. aryabhattai* were identified using bacterial genome sequencing (Supplemental Figure S1). The amino acid sequence of each identified endolysin was deduced from the DNA sequence, and its 3D structure was determined using Swiss-Model (42). The different domains present in the structure were determined using Pfam software (43). Alignment and phylogenetic trees were constructed based on the alignment of *B. altitudinis* and *B. aryabhattai* endolysin sequences with the amino acid sequences of bacteriophage endolysins available in the NCBI database. Protein sequences were aligned using Clustal X2 (44) and a phylogenetic tree was constructed using MEGA7 using the neighbor-joining method with P distance values (45). A phylogenetic neighbor-joining (NJ) tree of similarity was plotted using iTOL (version 7) (46) to classify them into different types.

### Analysis and molecular docking of the endolysins with peptidoglycan ligands

The quality of the predicted tertiary structures of the endolysins was assessed using the ERRAT, PROCHECK, and Verify 3D online tools (https://saves.mbi.ucla.edu). The binding pocket sites of endolysins were predicted using the P2Rank standalone method with the PrankWeb interface. The pocket with the highest score and probability was selected for docking analysis. The structures of the muramyl dipeptide and peptidoglycan monomers were obtained from the PubChem database (https://pubchem.ncbi.nlm.nih.gov/). Muramyl dipeptide (Compound CID: 451714) and peptidoglycan monomer (Compound CID: 5462260) were used as ligands during docking analysis (AutoDock Vina) using the PrankWeb interface. The endolysins with docked ligands and the distances in intermolecular interactions were visualized using PyMOL Molecular Graphics System (Version 1.2r3pre, Schrödinger, LLC). Further analyses and molecular docking were performed using ArtE2. Structural information for ArtE2 was obtained using PDBsum (47). The quality of conservation of amino acid residues of ArtE2 was determined using ConSurf (48).

### Recombinant expression and purification

The endolysins and ArtE2 encoding genes were amplified by PCR (Supplemental Figures S1 and S2) (Table 1), digested with NdeI/BamHI restriction enzymes, and ligated with T4 ligase (NEB) into the same restriction sites in pCold I (Takara, Japan), resulting in pCold I-ArtE2 encoding a lysin with a C-terminal hexa-histidine tag. The clones were confirmed using Sanger sequencing (Sangon Biotech, China). The recombinant expression of different endolysins (E1, E2, E3, E7, E10, E12, and E15) and engineered endolysin 2 (Art2) was performed in *E. coli* BL21(DE3). In the case of the construction of the coding gene of Art2, it was designed based on the sequence of the wild type endolysin E2 (NCBI accession code PQ842537). A polycationic nanopeptide (amino acid sequence: KRKKRKKRK) was fused to the N-terminal end of endolysin. Codon optimization of the gene sequences was performed. The gene coding sequences of the endolysins and ArtE2 were cloned into the pCold I vector. The recombinant plasmids were transformed into *E. coli* BL21(DE3) cells (Sangon Biotech, Shanghai, China) and verified by sequencing.

**TABLE 1.**
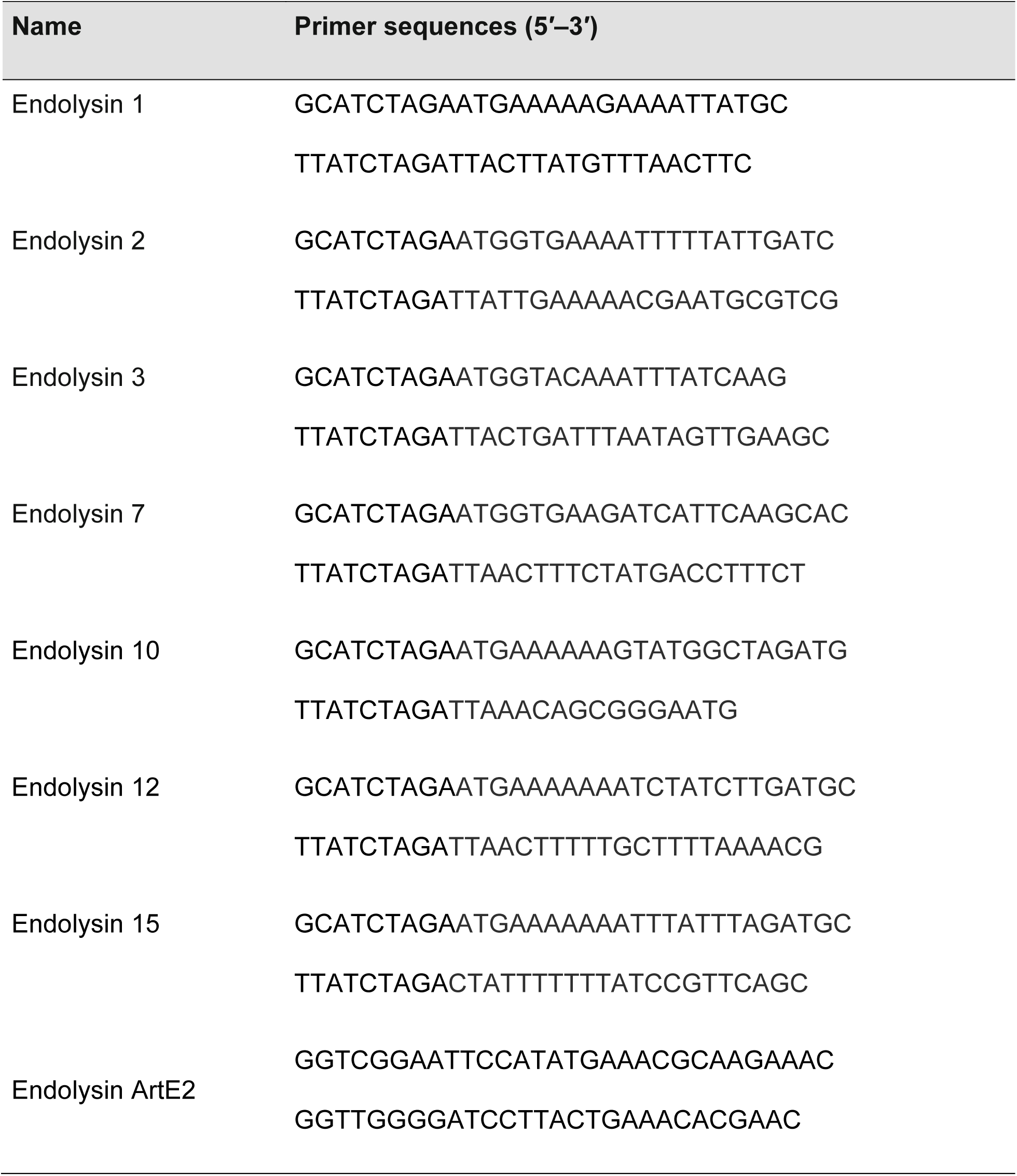
List of primers used in this work.

After screening on LB agar plates with 100 μg/mL of ampicillin to obtain single recombinant colonies, the obtained single recombinant colonies were inoculated in LB broth medium containing 100 μg/mL of ampicillin and cultured at 37°C (150 rpm) until the optical density (OD) at 600 nm reached 0.6. Subsequently, IPTG was added at a final concentration of 0.5 mM, followed by incubation for 18 h at 15°C. The cells were then collected by centrifugation at 4000 × g for 40 min and resuspended in 25 ml of lysis buffer (300 mM NaCl, 20 mM Tris-HCl (pH 7.5), and 1% Triton X-100 (v/v)). Cell disruption was performed by sonication (Goldenwall, China) under the following conditions: 45% maximum amplitude (120 watts) for 25 min with 30-s pulse/30-s break steps on ice. The lysate sample was centrifuged at 10,000 rpm for 30 min, and the supernatant was passed through a 0.22 μm PES membrane (Tianjin Jinteng Experiment Equipment Co., Ltd., China). ArtE2 was purified using a HisPur Ni-NTA Gravity Column (Thermo Fisher Scientific, US) according to the manufacturer’s instructions. The identity and purity of ArtE2 were verified by sodium dodecyl sulfate-polyacrylamide gel electrophoresis (SDS-PAGE).

### Expression analysis by Western blotting

Western blotting was performed to confirm the presence of the expected amount of the ArtE2 protein in the eluate. Proteins were resolved by SDS-PAGE and transferred to a polyvinylidene fluoride (PVDF) membrane (Biosharp, China) using a semi-dry transfer method (Sangon Biotech, China). After blocking the membrane for 1 h with 5% (w/v) skimmed milk in PBS (milk/PBS) (Labcoms, US), it was incubated overnight at 4 °C with a 1:1000 dilution of the primary antibody: mouse serum containing anti-6xHis antibody (Sangon Biotech, China). After three washes with Tris-buffered saline containing 0.1% Tween 20, membranes were incubated with a 1:10,000 dilution of goat anti-mouse horseradish peroxidase (HRP) secondary antibody (Sangon Biotech) for 1 h at room temperature. After three washes with TBST (Sangon Biotech, China), the blot was incubated in a 1:1 ratio with Western Bright ECL solution (Thermo Fisher Scientific, US). The ChemiDoc MP Imaging System (Liuyi Biotechnology, China) was used to capture ECL signals.

### Antibacterial activity

The biological activities of endolysins (E1, E2, E3, E7, E10, E12, and E15) were evaluated using the inhibition zone method. *Staphylococcus aureus* was grown in liquid medium (LB broth) until it reached the exponential phase (OD600 = 0.6). The culture was diluted to 10 CFU/mL to standardize its concentration. A total of 100 µL of the standardized bacterial culture was evenly spread across the surface of an agar plate using a sterile spreader. Sterile filter paper discs (6 mm diameter) were soaked in 20 µL of each endolysin solution at 2 µg/mL and placed on the inoculated agar. Buffers without endolysins or water were used as the controls. The plates were incubated at 37°C for 24 h. The diameter of the inhibition zone was measured in millimeters.

### Estimation of minimum inhibitory concentration

*Staphylococcus aureus* and *Escherichia coli* in the mid-exponential growth phase (OD600 = 0.6) were harvested by centrifugation (16,000 × g for 5 min), washed, and diluted 100-fold in 10 mM and 5 mM HEPES (pH 7.4) to a final density of 10^8 CFU/ml, respectively. A protein solution containing ArtE2 (0, 2.31, 5.21, 6.95, and 8.11 µg/mL) in 10 mM HEPES (pH 7.4) was added to 50 μl of the S. aureus cell dilution in a final volume of 100 µL and incubated at 37 °C with shaking at 100 rpm. After 60 min of incubation, dilutions of the cell suspensions were plated on LB agar plates in triplicates. colonies were counted after overnight incubation at 37 °C. Meanwhile, a solution containing ArtE2 (0, 2.31, and 5.21 µg/mL) in 5 mM HEPES (pH 7.4) was added to 50 μl of the E. coli cell dilution in a final volume of 100 µL and incubated at 37 °C, with shaking at 100 rpm. After 30 min of incubation, dilutions of the cell suspensions were cultivated in LB broth in triplicates. OD600 was evaluated after overnight incubation at 37 °C. LB medium and water were used as the controls. Antibacterial activity was determined by linear correlation analysis using the Pearson’s r test.

### Time-kill curve

Mid-exponentially growing S. aureus and E. coli cells (OD600 = 0.6) of *S. aureus* and *E. coli* in LB broth were harvested by centrifugation (16,000 × g for 5 min), washed, and diluted 100-fold in 10 mM and 5 mM HEPES (pH 7.4) to a final density of 10^8 CFU/ml, respectively. A suspension with a total volume of 100 μl containing 50 μl of bacterial cell dilution and 50 μl of ArtE2 (5.21 µg/mL) in 10 mM ( *S. aureus*) and 5 mM ( *E. coli*) HEPES (pH 7.4) was prepared. These mixtures were incubated at 37 °C in a shaker for the following times: a) for S. aureus at 0, 15, 30, 45, and 60 min; and b) for *E. coli* at 0, 10, 15, 20, and 25 min. The turbidity of the culture was monitored at the respective time points and the reduction was determined. Antibacterial activity was determined by linear correlation using the Pearson’s r test.

### Statistical analysis

All experiments were performed in triplicate, unless otherwise specified. Data are presented as mean ± standard deviation. One-way ANOVA followed by Tukey’s post hoc analysis was used to determine the significance of the treatments. Differences between the means were considered significant at P < 0.05 and highly significant at P < 0.01. Antibacterial activity was determined by linear correlation using the Pearson’s r test. Data were analyzed and processed using GraphPad Prism software (La Jolla, CA, USA).

## RESULTS

### Isolation and characterization of Bacillus endolysins

Seven different endolysin genes (E1, E2, E3, E7, E10, E12, and E15) were discovered and isolated from Bacillus altitudinis and Bacillus aryabhattai phages using deep sequencing (Dataset D1; GenBank accession number: PRJNA781678). Structural domain analysis showed that each of the seven endolysins contained a conserved enzymatically active domain (EAD) and N-acetylmuramoyl-L-alanine amidase domain (Figure 1A). In terms of the structure of their cell wall-binding domains (CBDs), endolysin-2 has an SPOR domain, endolysin-3 and endolysin-12 have two different LysM and peptidoglycan-binding domains, endolysin-7 and endolysin-10 have peptidoglycan-binding and LysM domains, endolysin-15 has a PlyG domain, and endolysin-1 has a CBD with an unknown structure (Figure 1A).

**FIG 1.**
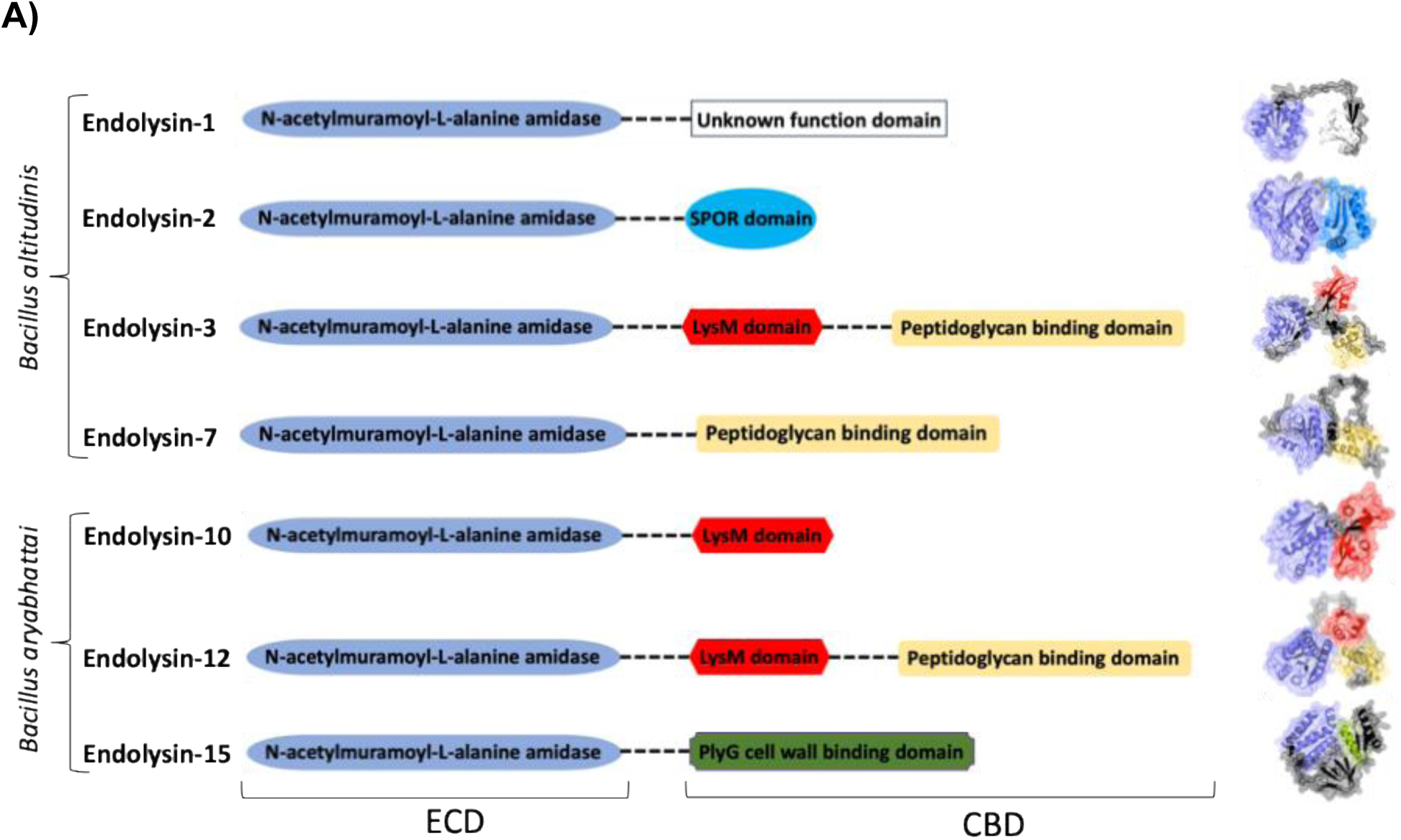
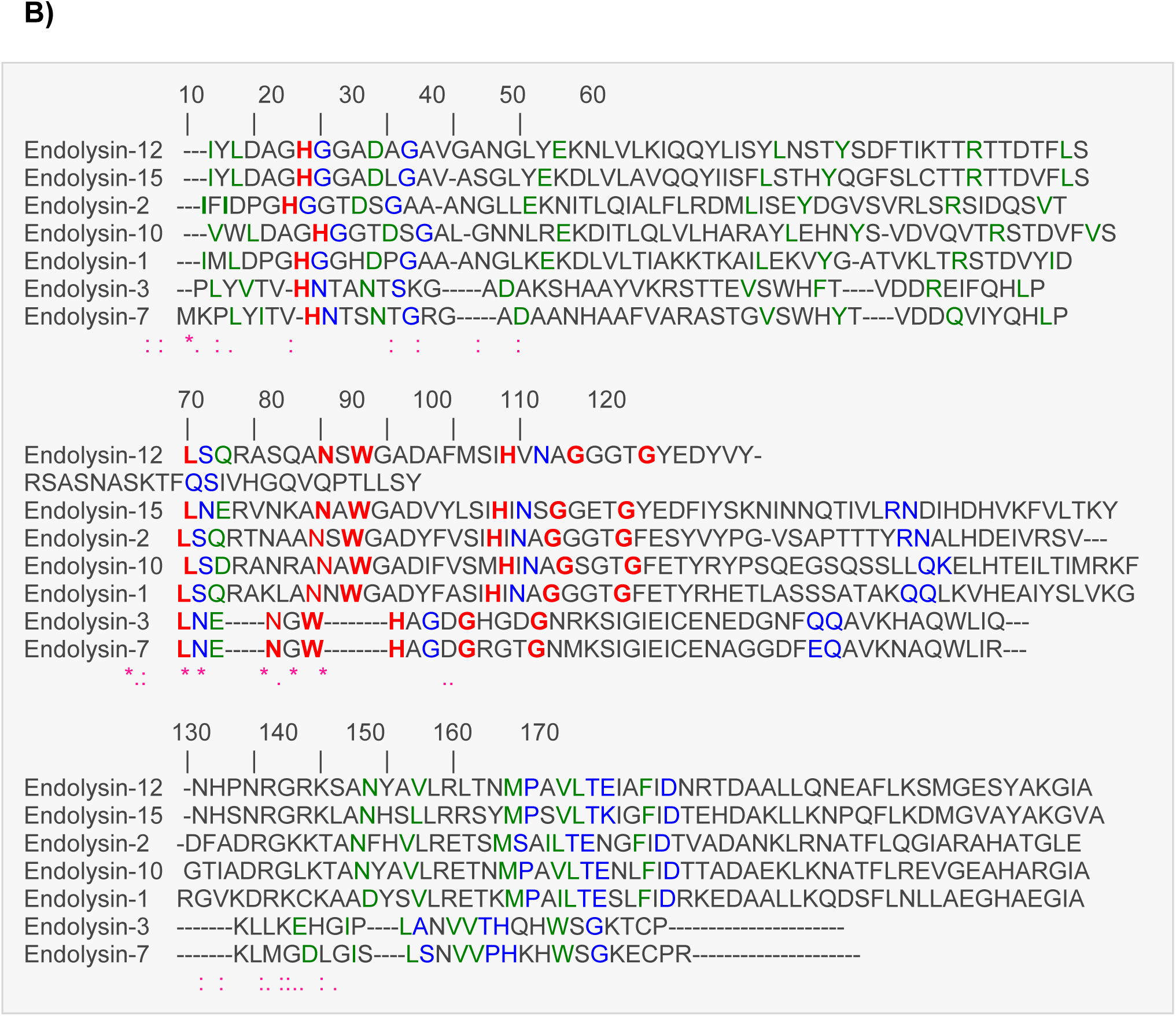
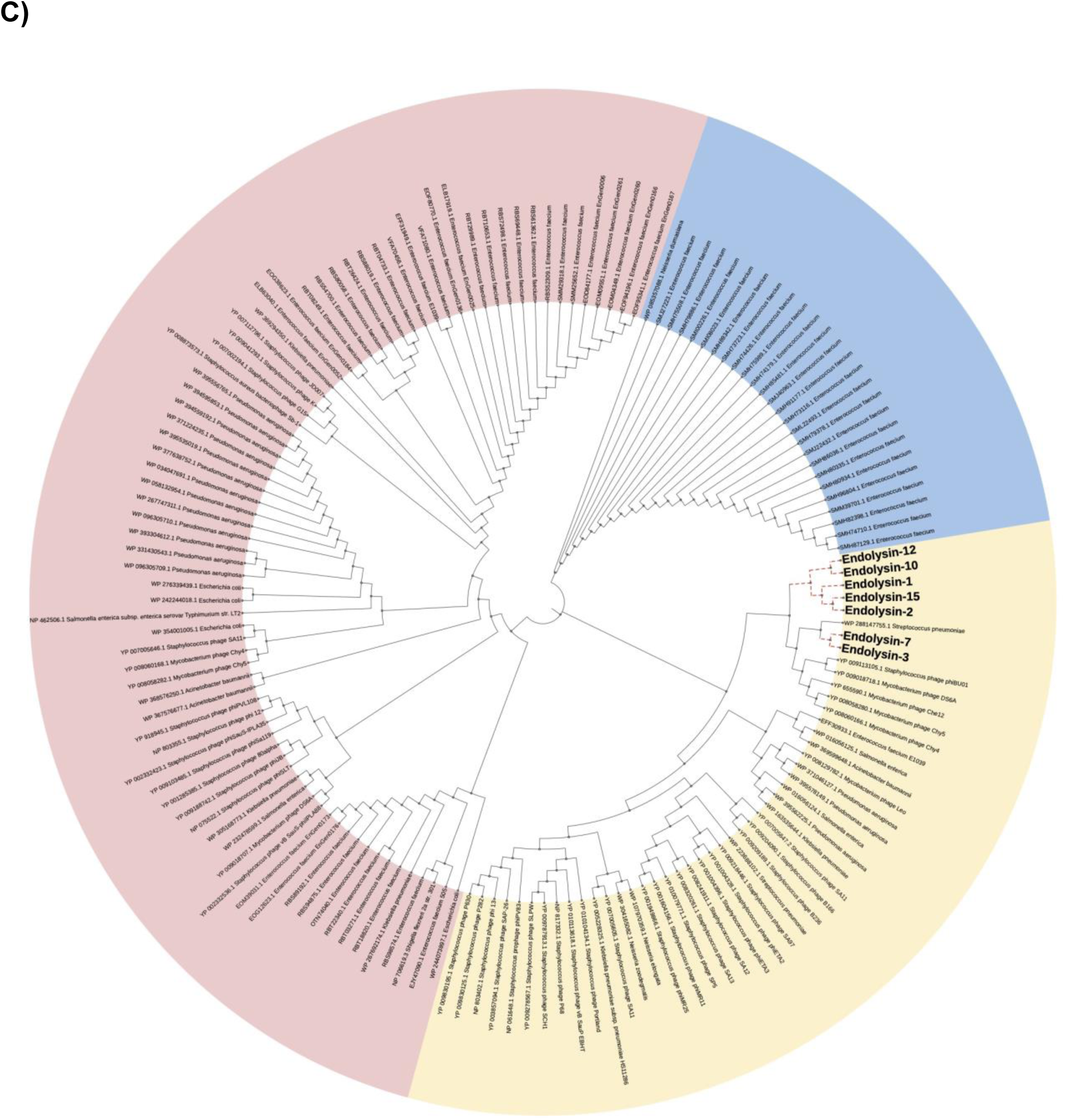
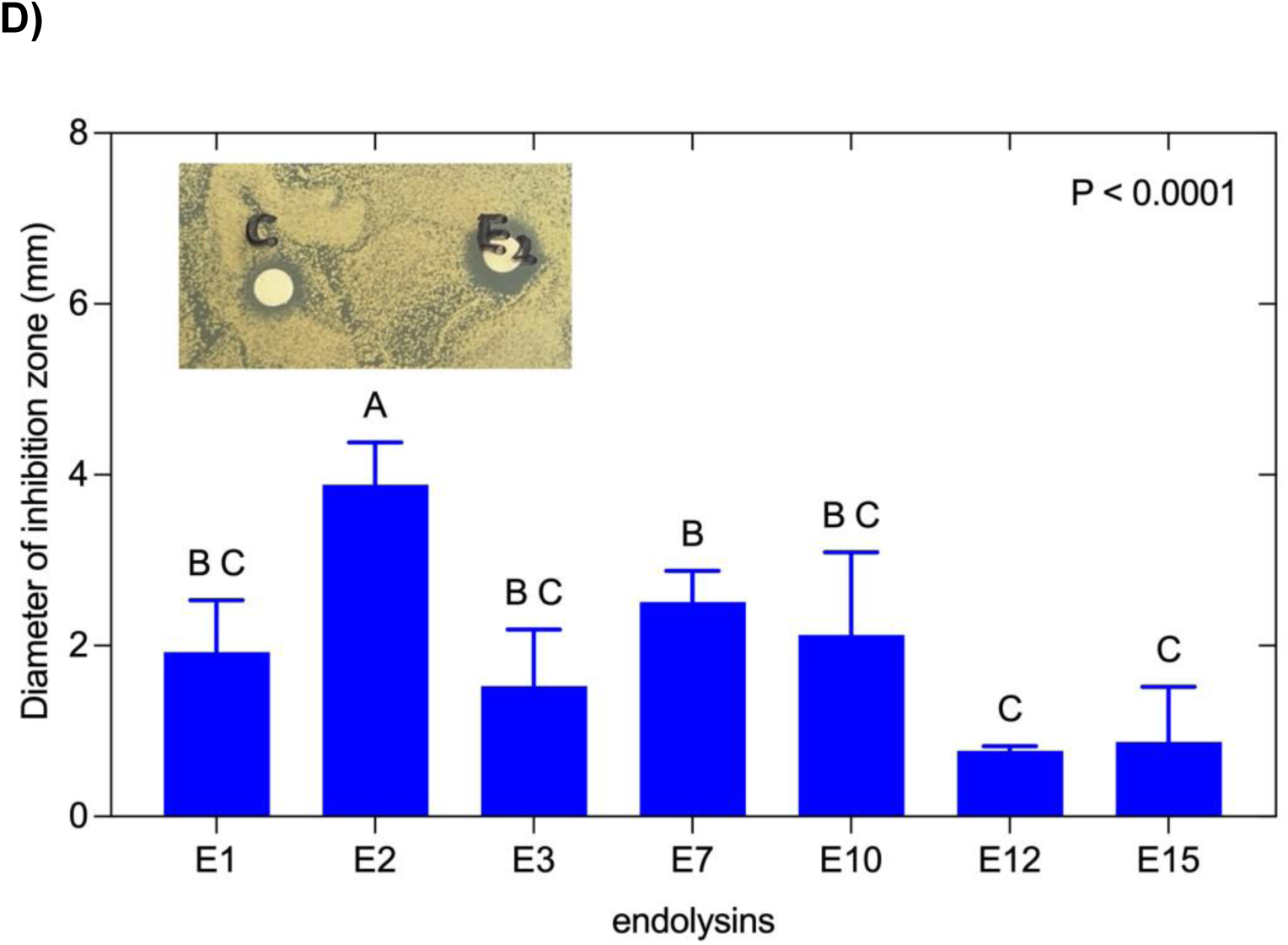
*In silico* and comparative analysis of endolysins isolated from *Bacillus altitudinis* and *Bacillus aryabhattai* species. **A)** Diversity of enzymatic catalytic domain (ECD), cell-binding domain (CBD), and deduced three-dimensional models found in *B. altitudinis* and *B. aryabhattai* endolysins. **B)** Amino acid sequence alignment of enzymatic catalytic domains of *B. altitudinis* and *B. aryabhattai* endolysins. Alignments were performed using Clustal Omega. Conserved amino acids are shown in red bold font with asterisks. (**C)** Phylogenetic relationship between *B. altitudinis* and *B. aryabhattai* endolysins and other endolysins from different bacterial species. Neighbor-joining trees were constructed based on amino acid sequences using MEGA X with 1000 bootstrap replicates and the iTOL program. **D)** Lytic activity assay of endolysin extracts against *Staphylococcus aureus*. Filter papers were soaked in crude extract suspended in 20 mM Tris-HCl (pH 8.0) of different endolysins from recombinant *E. coli* BL21(DE3) and *E. coli* BL21(DE3) containing pCold I vector empty (control) and placed onto a bacterial lawn of *S. aureus*. The area of inhibition was measured in millimeters. The bars represent the means of three independent replicates ± standard error (P < 0.0001).

Multiple sequence alignment of the catalytic domains demonstrated significant conservation of residues, such as histidine (H), leucine (L), asparagine (N), tryptophan (W), and glycine (G), especially around amino acid positions 10-20, 40-50, and 90-100 (Figure 1B). Less conservation and higher sequence variability occurred near positions 60-80 and 120-130. Variable C-terminal regions of endolysins-3 and -7 were observed, demonstrating gaps from positions to 120-170, while the N-terminus (positions 1-50) was generally less conserved across all sequences (Dataset D2).

Phylogenetic analysis using neighbor-joining trees grouped Bacillus endolysins within a clade containing 48 endolysins from diverse bacterial genera, including Streptococcus, Staphylococcus, Mycobacterium, Enterococcus, Salmonella, Acinetobacter, Pseudomonas, Klebsiella, and Neisseria (Figure 1C). Notably, endolysins-3 and -7 clustered closely with Streptococcus pneumoniae endolysins, indicating a potentially shared specificity or evolutionary relationship.

Biological activity assays assessing lytic capacity against Staphylococcus aureus BM9393 revealed that endolysin-2 exhibited the highest activity, producing inhibition zones approximately 5 mm in diameter (Figure 1D). Endolysins-7 and -10 also showed significant lytic activity, with inhibition zones of approximately 3 mm, whereas endolysins-1, -3, -12, and -15 demonstrated smaller inhibition zones, ranging from 1 to 2 mm. The empty vector control did not exhibit inhibition, confirming the enzymatic specificity. These results highlight endolysin-2 as the most promising candidate for further studies.

### In silico analysis of endolysin ArtE2

Using a combination of computational methods, we predicted the three-dimensional structure of ArtE2, an artificially created endolysin-2 fused with the polycationic peptide KRKKRKKRK at the N-terminus, to identify the structural elements that determine its function. The Pfam and Conserved Domain Database (CDD) assigns the catalytic domain to the first 168 amino acids (residues 4-171) at the N-terminus of the protein, whereas the cell wall-binding domain is located at the C-terminus (residues 181-250) (Figure 2A).

**FIG 2.**
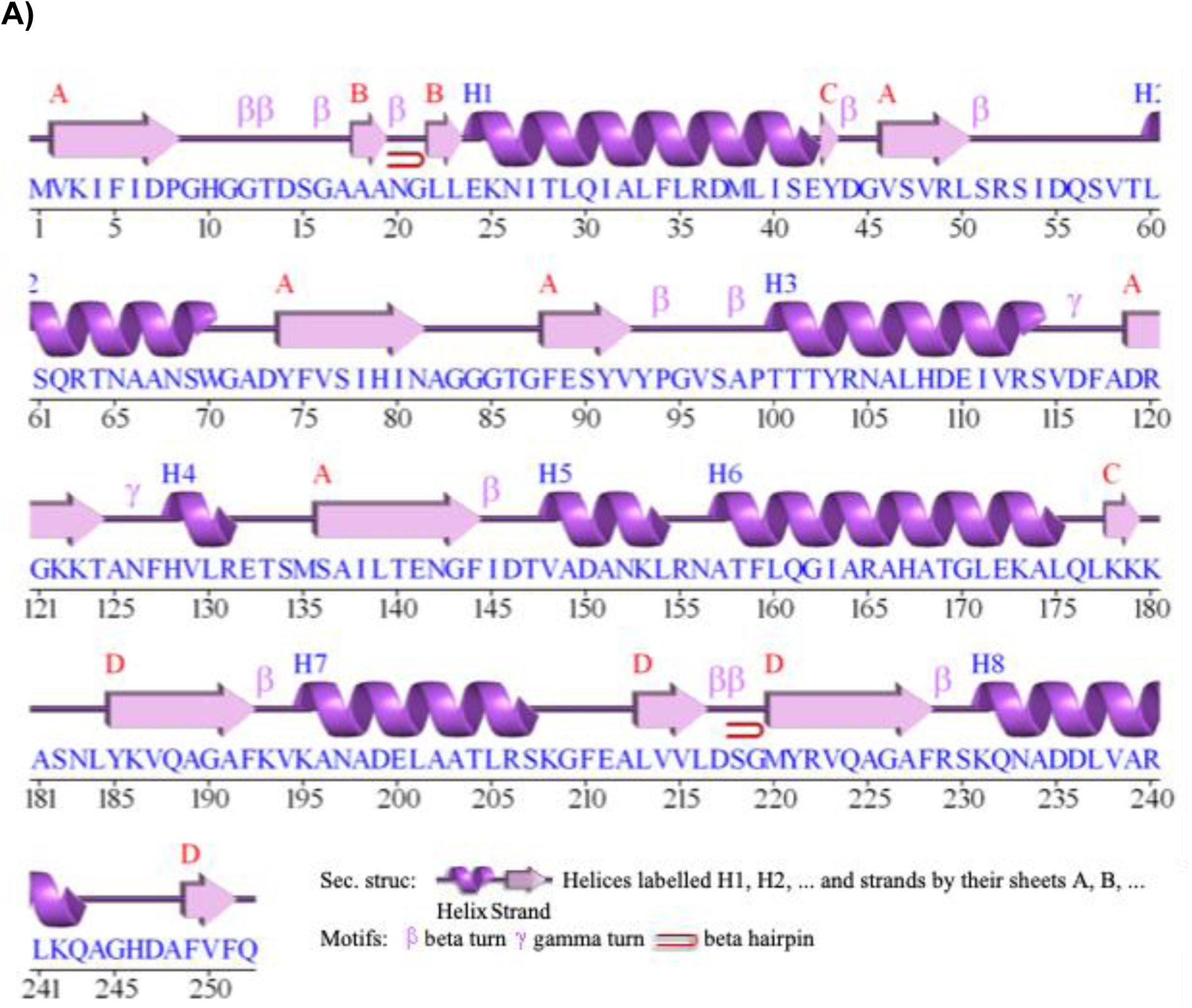
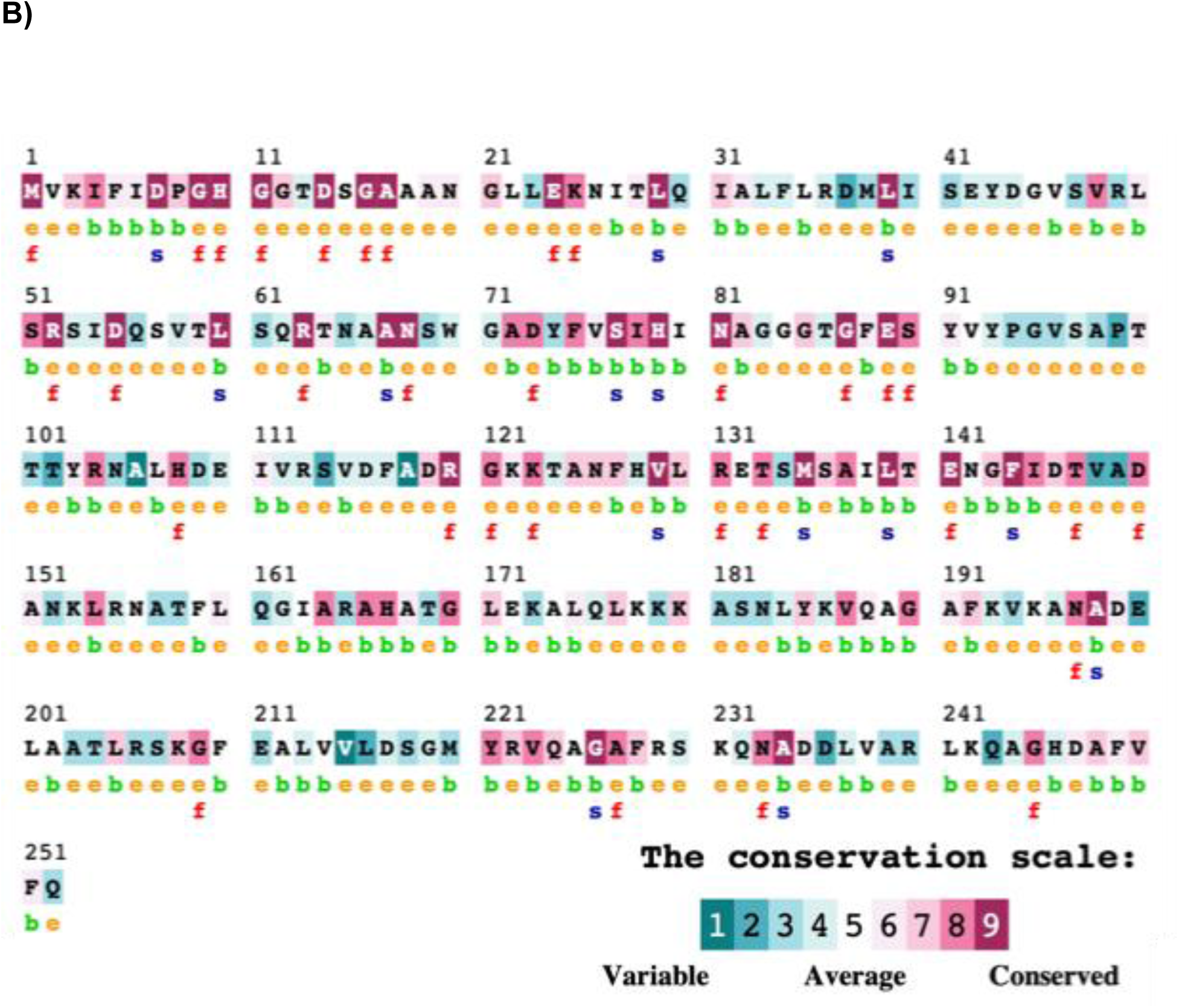
*In silico* characterization of the endolysin ArtE2. **A)** Predicted secondary structure of ArtE2 built using PDBsum. **B)** Conservation quality of amino acid residues of ArtE2 as determined by ConSurf. This indicates the distribution of structural and functional residues in the structure. Conservation scores of amino acids were on a scale varying from 0 to 9, indicating variable to conserved amino acids. e: an exposed residue according to the neural network algorithm; b: a buried residue according to a similar algorithm; f: predicted functional residue (highly conserved and exposed); s: predicted structural residue (highly conserved and buried).

PDBsum secondary structure prediction identified α-helical structures throughout the sequence, specifically at residues 25–35, 70–80, 100–110, 150–160, 170–180, 200–210, 220–230, and 240–250, and β-sheet structures at residues 40–50, 60–70, 130–140, 190–200, 220–230, and 240–250 ( Figure 2A). Beta turns, gamma turns, and beta hairpin motifs were the most abundant in regions 1 and 4. These flexible structural elements may be involved in the region-specific structural transitions required for substrate recognition.

ConSurf conservation mapping revealed that 30 of the 246 amino acid positions were conserved among all E2 endolysins, with 16 being both surface-exposed and functionally conserved. These include Gly-9, His-10, Asp-14, Glu-24, Arg-63, and Glu-89, which are essential for catalytic or substrate binding reactions (Figure 2B). Fourteen additional positions were also found to be highly conserved but internalized within the protein and therefore likely contribute to the structural stability of the molecule.

### Molecular docking of endolysin ArtE2

Ligand-binding pocket prediction using PrankWeb revealed two possible binding sites for ArtE2 (figure 3A). Pocket 1 has a much higher overall score (32.44) than Pocket 2 (4.02), and is therefore given a much greater probability (.903) of being functionally relevant. The pocket contained many highly conserved and accessible residues such as Gly-9, His-10, Gly-11, Asp-14, Ser-15, Gly-16, Ala-17, Glu-24, Arg-63, Asn-81, Glu-89, Leu-139, and Glu-141. Therefore, it is highly likely that this is the active site of ArtE2.

**FIG 3.**
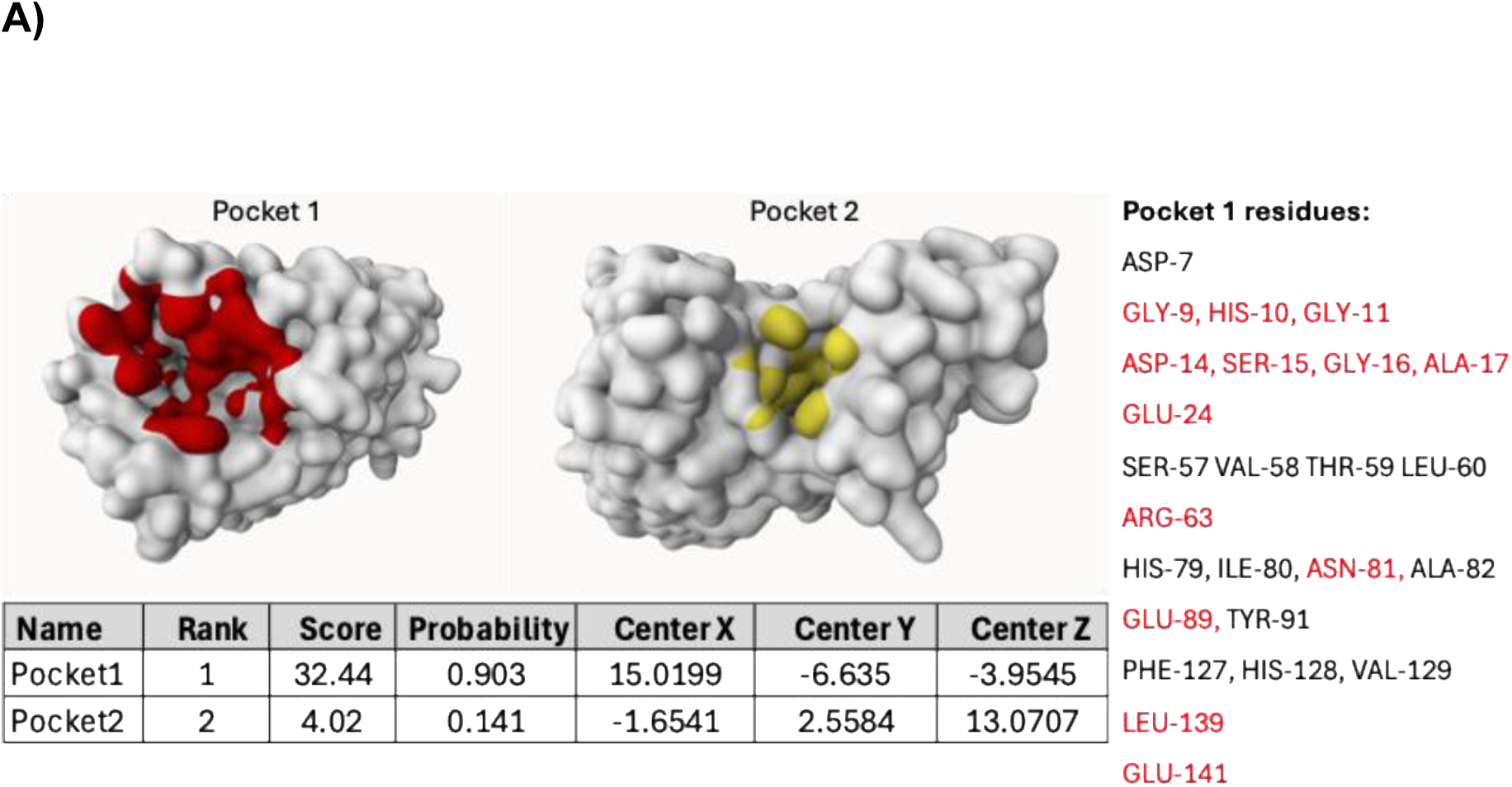
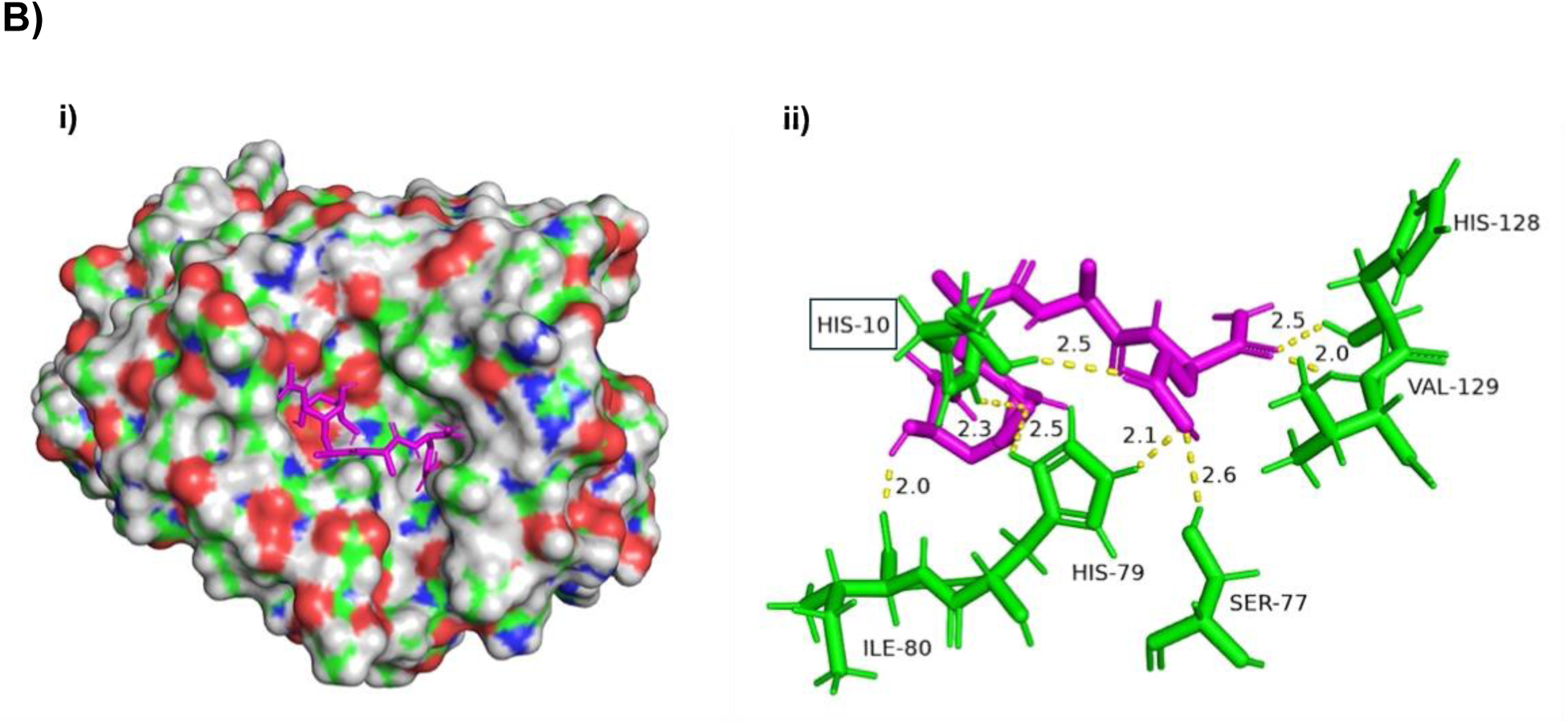
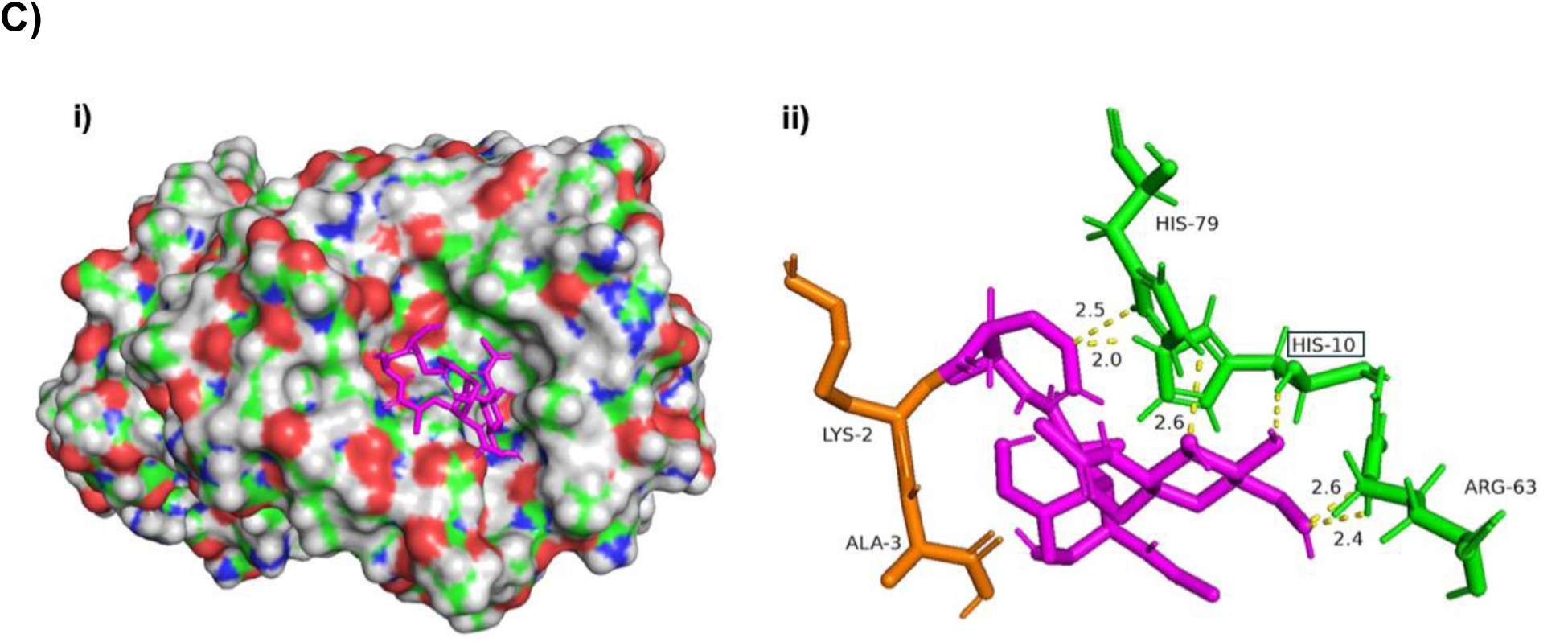
Analysis of docking interactions between endolysin ArtE2 and peptidoglycan. **A)** Prediction of ligand-binding sites on the surface of endolysin ArtE2 using the PrankWeb tool (https://prankweb.cz/). The ligand-binding pockets of ArtE2 are shown in red (pocket 1) and yellow (pocket 2). The figure also shows the amino acids in pocket 1 with a higher score (32.98) and the amino acids that are highly conserved and exposed in the endolysin. The panel below summarizes the binding sites of endolysins. **B)** Structure of ArtE2 in complex with muramyl dipeptide. **i)** Molecular surfaces of ArtE2 domains are colored according to their electrostatic surface potential generated by APBS (±5 kT/e): red, negatively charged; white, neutral; and blue, positively charged regions. **ii)** Visualization of the interaction between the active site residues (green) and muramyl dipeptide (magenta) with the distances for the hydrolysis of the peptidoglycan. Putative key catalytic residues are shown as rectangles. **C)** Structure of ArtE2 interacting with peptidoglycan monomer. **i)** Molecular surfaces of ArtE2 domains are colored according to their electrostatic surface potential generated by APBS (±5 kT/e): red, negatively charged; white, neutral; and blue, positively charged regions. **ii))** Visualization of the interaction between the active site residues (green) and peptidoglycan monomer (magenta) with the distances for the hydrolysis of the peptidoglycan. Putative key catalytic residues are shown in a rectangle. AlphaFold 2.0 was used to generate high-confidence models of endolysins, and docking analysis was performed using PrankWeb. The visualization was based on a PyMOL script (http://www.pymol.org/pymol).

Molecular docking studies using AutoDock Vina were performed to assess the interactions between ArtE2 and both muramyl dipeptide (MDP) and a peptidoglycan monomer, which are important structural components of the bacterial cell wall (figures 3B and 3C). The docking scores were -6.5 (MDP) and -5.9 (peptidoglycan monomer), indicating that ArtE2 binds to these substrates in an energetically favorable manner (Table 2).

**TABLE 2.** Ligand binding site prediction and docking score during the interaction between the endolysin ArtE2 and muramyl dipeptide/peptidoglycan monomer.

|  | Ligand Binding Site |  | Docking Analysis |  |
| --- | --- | --- | --- | --- |
|  | Prediction |  | Muramyl | Peptidoglycan |
|  | (PrankWeb) |  | Dipeptide | Monomer |
|  | Score | Probability | Score | Score |
| ArtE2 | 32.44 | 0.932 | -6.5 | -5.9 |

A detailed analysis of the individual residue interactions demonstrated that several histidine residues (His-10, His-79, and His-128) acted either catalytically or stabilized the substrate. His-10 forms hydrogen bonds with MDP at 2.3 Å and 2.5 Å distances, whereas His-79 and His-128 also form hydrogen bonds at 2.3 Å and 2.5 Å distances, respectively (figure 3B(ii)). Arginine-63 is involved in ionic and hydrogen bond interactions with the peptidoglycan monomer at 2.4 Å and 2.6 Å distances, respectively, and demonstrates how the substrate is positioned and bound (figure 3C(ii)). Other residues, such as Ser-77, Ile-80, and Val-129, contribute to the substrate via hydrophobic contact and hydrogen bonding to accommodate the ligand. Surface electrostatic mapping of ArtE2’s active site shows that the area is composed of both positively and negatively charged areas, which will help in the recognition and catalysis of the substrate.

### Recombinant expression of ArtE2 endolysin

ArtE2 was cloned into the pCold I vector and expressed in Escherichia coli BL21(DE3). SDS-PAGE analysis confirmed the overexpression of a 28 kDa protein corresponding to ArtE2 (Figure 4A). Purification by Ni-NTA affinity chromatography resulted in >90% pure protein, as evidenced by a sharp distinct band at the expected molecular weight. Western blotting with an anti-6xHis antibody validated the identity of recombinant ArtE2, detecting bands in purified fractions consistent with SDS-PAGE results (Figure 4B), confirming successful expression and purification.

**FIG 4.**
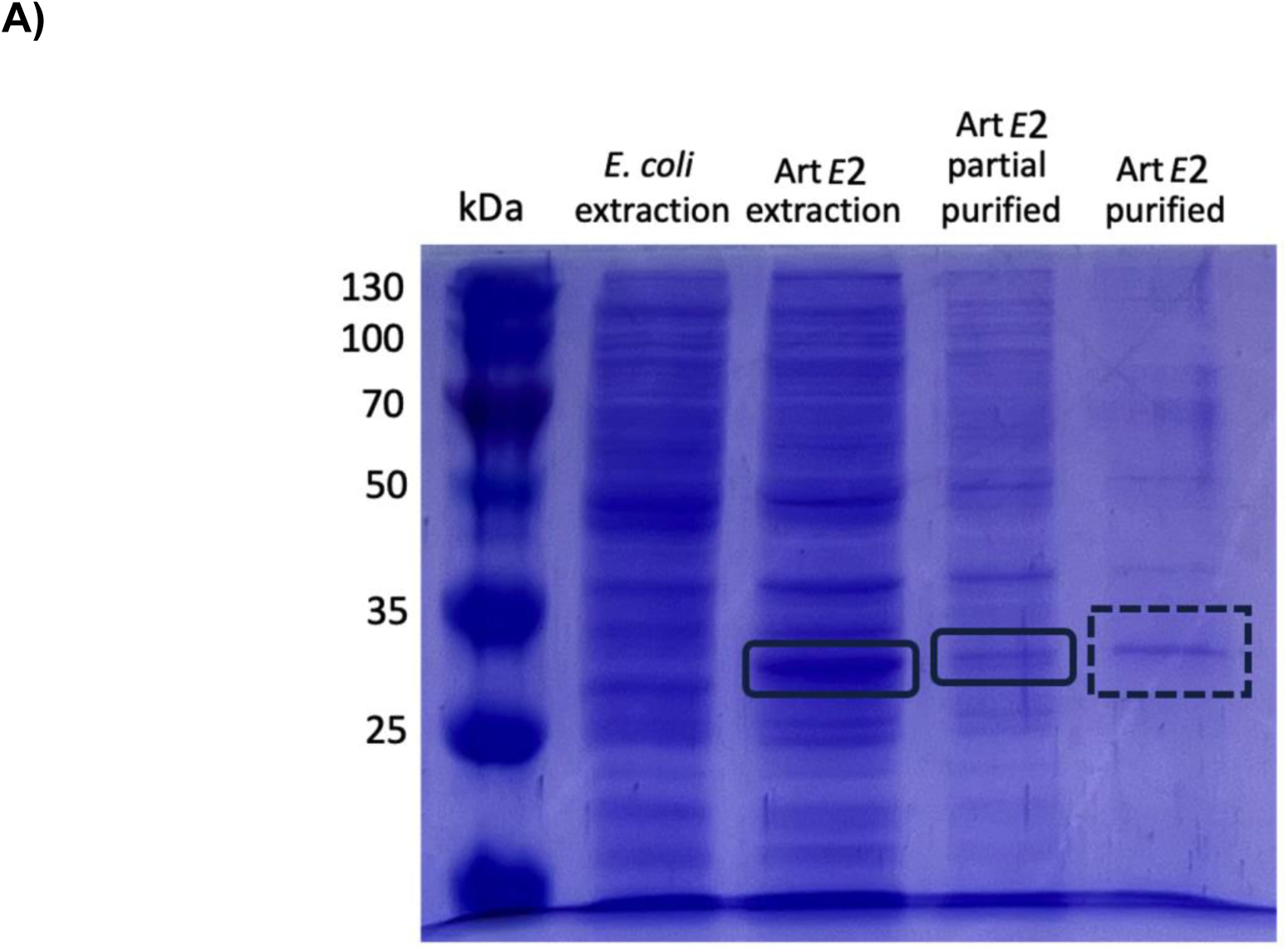
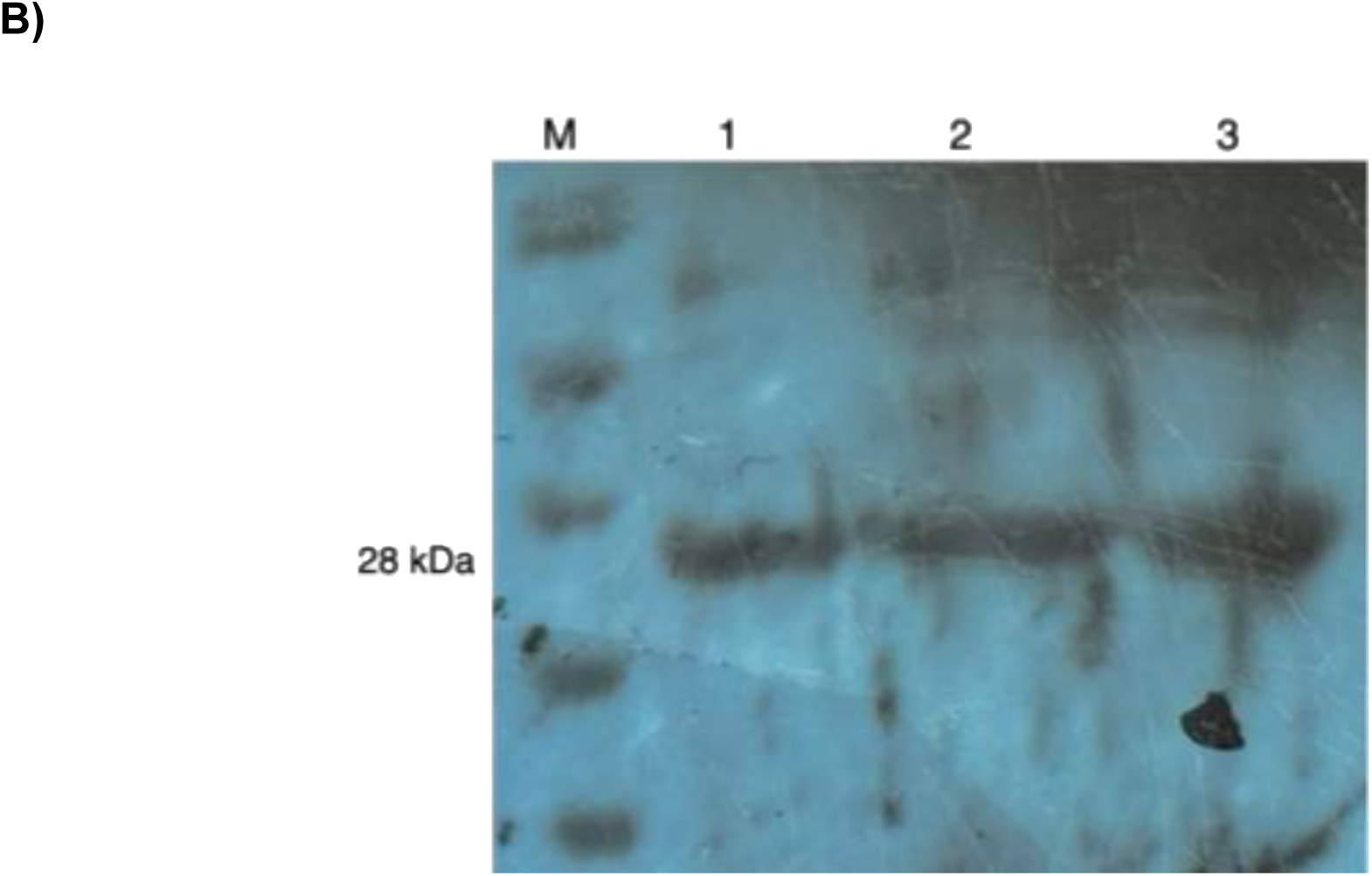
Analysis of recombinant endolysin ArtE2 expressed in *Escherichia coli* BL21(DE3). **A)** SDS-PAGE for ArtE2 production analysis. Purified pCold I::ArtE2 was loaded onto an SDS-PAGE gel. Lane 1, molecular mass marker (SAGON BIOTECH); lane 2, E. coli extract; lane 2, ArtE2 extract; lane 3, ArtE2 partially purified; and lane 4, ArtE2 purified. **B)** Western blot analysis of recombinant ArtE2 from *E. coli* BL21(DE3) (pCold I::ArtE2) using an anti-His tag antibody. M: Molecular mass marker (SAGON BIOTECH). Lanes 1, 2, and 3 show *E. coli* BL21(DE3) (pCold I::ArtE2).

### Biological activity of ArtE2 endolysin

For *Staphylococcus aureus* BM9393, dose-dependent killing was indicated by the presence of viable colonies. As shown in Figure 5A, as ArtE2 concentration increased from 0 µg/mL to 8.11 µg/mL, the number of viable colony-forming units (CFU/mL) per mL decreased until it was reduced to almost zero for all colonies. This was consistent with the ability of the enzyme to kill bacteria in a dose-dependent manner. A Pearson correlation coefficient of r = -0.538 was observed between the enzyme concentration and percentage of surviving bacteria. Additionally, time-kill assays were performed on S. aureus BM9393 cells for 60 minutes. As shown in Figure 5B, there was a significant decrease in the optical density (OD) after 15 min, and this trend continued until 60 min post-treatment. These data suggest that the enzyme causes rapid lysis of bacterial cells. Furthermore, visual inspection of the plates indicated a decline in bacterial cell density, which was correlated with the rapid killing activity of the enzyme.

**FIG 5.**
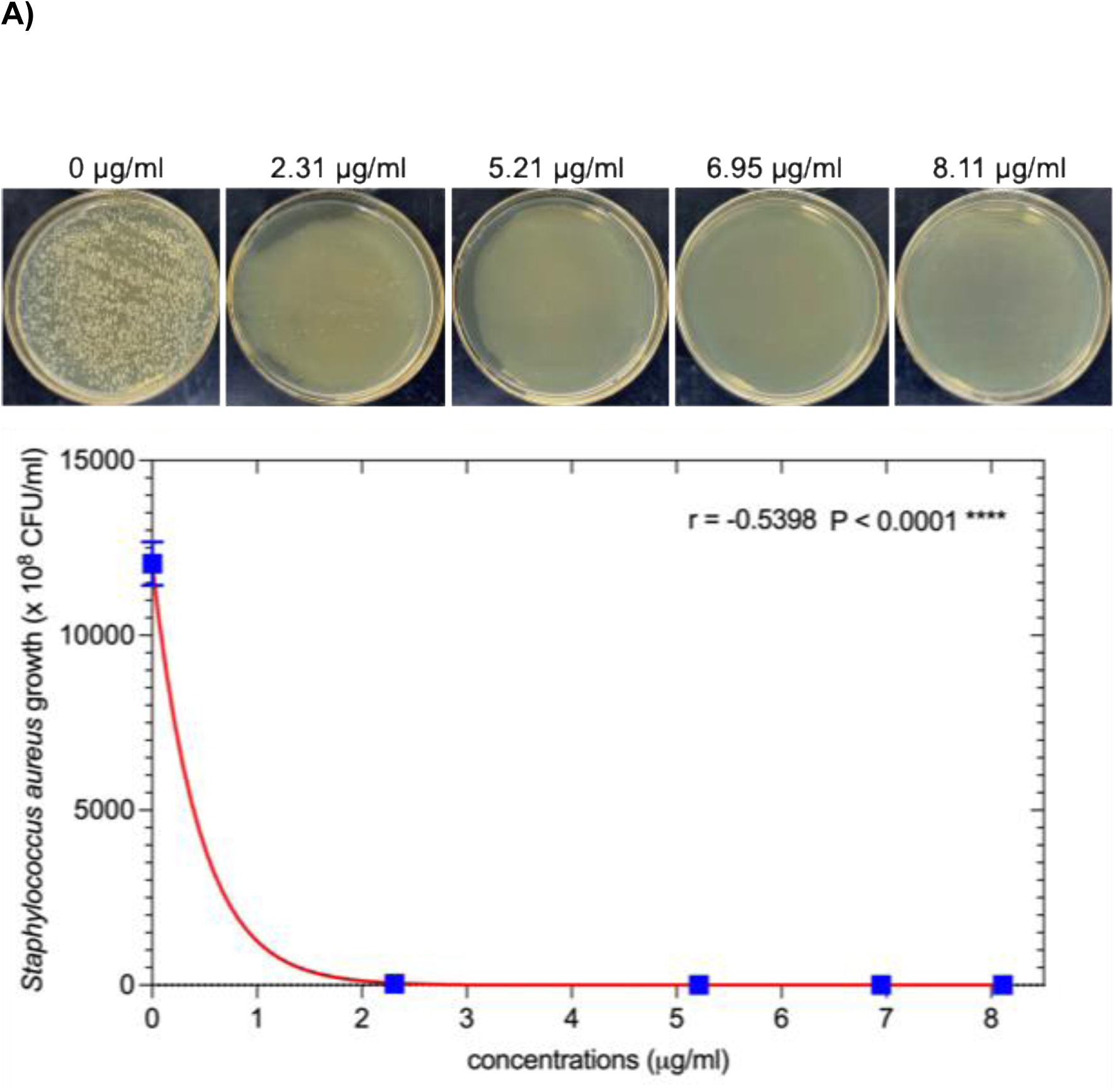
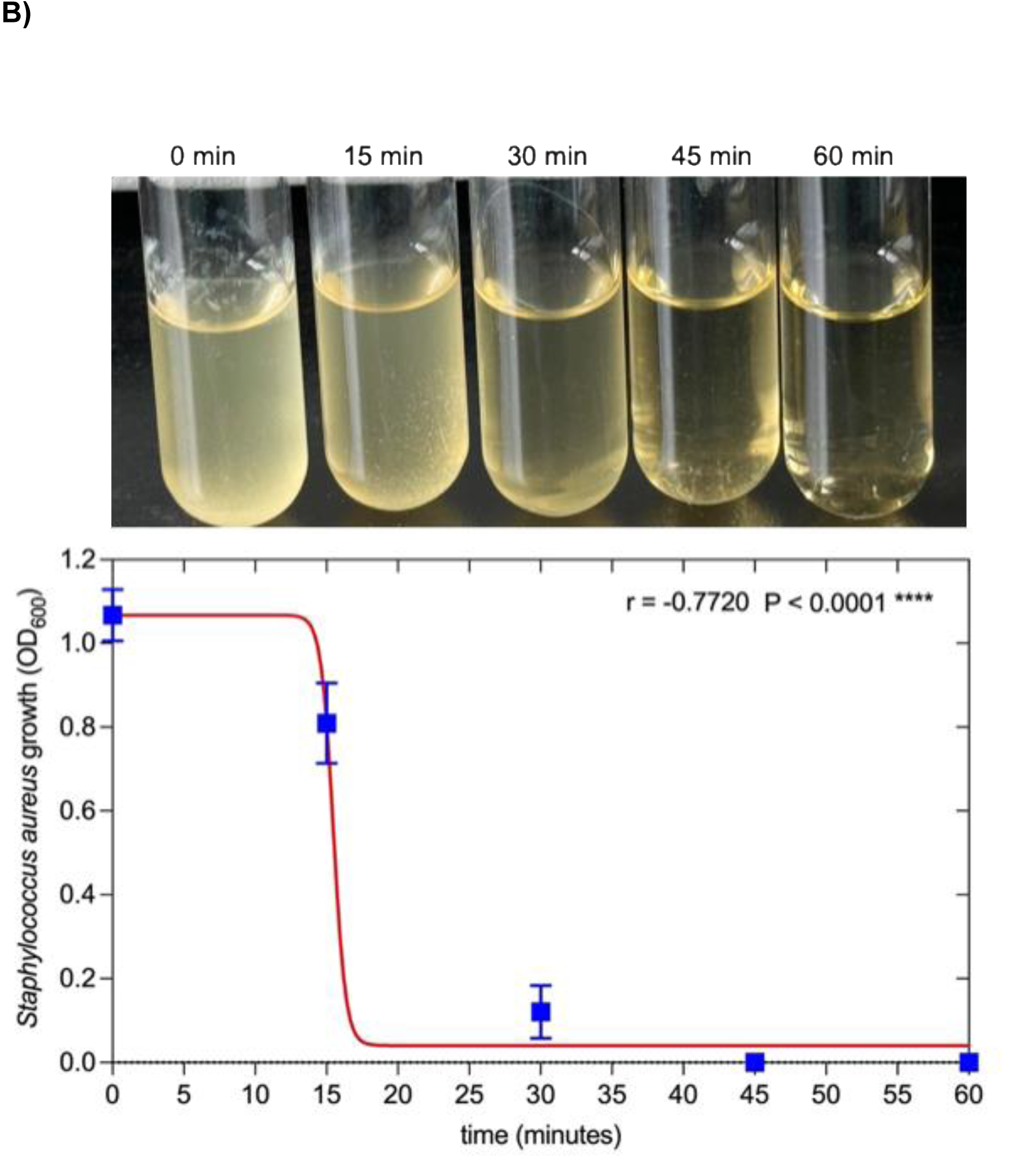
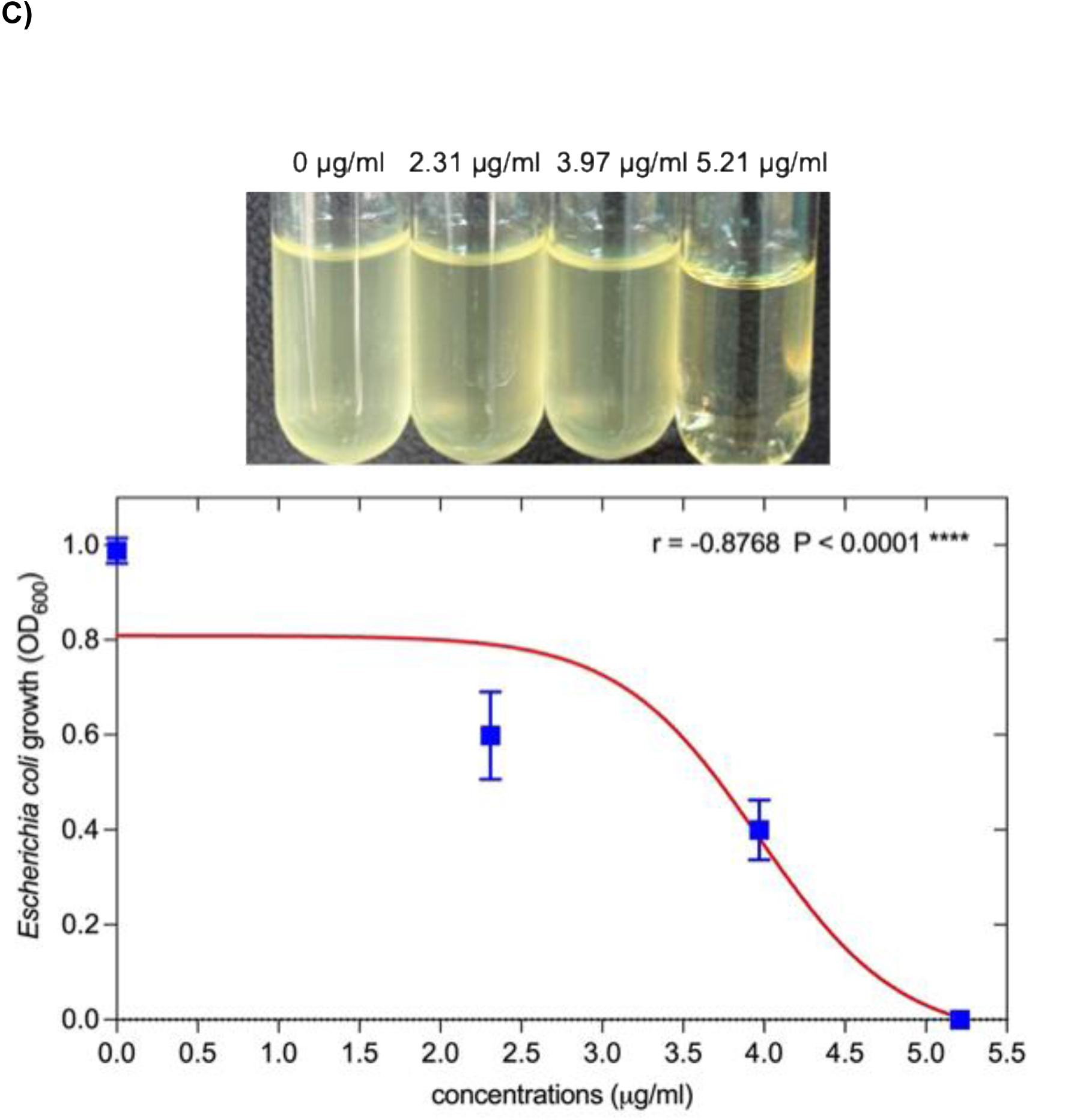
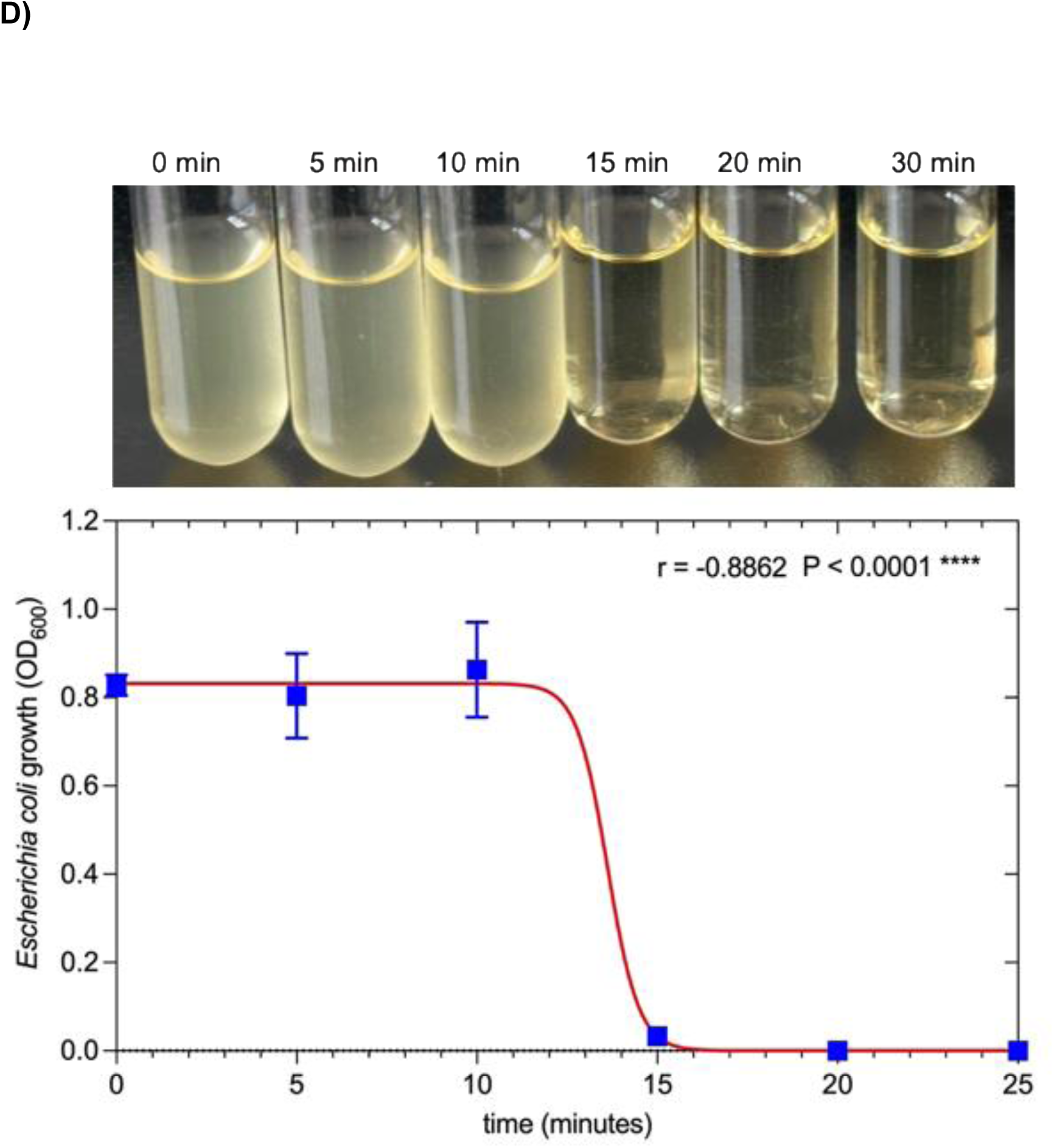
Antibacterial activity of endolysin ArtE2 against *Staphylococcus aureus* BM9393 and *Escherichia coli* BL21(DE3) cells. **A)** Different concentrations of endolysin were added to washed *S. aureus* BM9393 cells resuspended in incubation buffer (20 mM Tris-HCl, pH 8.0). The number of viable colonies was counted after treatment. **B)** Time-kill experiment of *S. aureus* BM9393 treated with ArtE2 at pH 7.4. (**C)** Activity of ArtE2 in *E. coli* BL21(DE3) at different concentrations. Enzyme activity was measured spectrophotometrically at 600 nm by measuring the changes in turbidity. **D)** Evaluation of the treatment of *E. coli* BL21(DE3) at different time points with ArtE2.

In the case of *Escherichia coli* BL21(DE3), the growth of *E. coli* was inhibited in a dose-dependent manner by ArtE2. As illustrated in Figure 5C, the OD600 values decreased as the enzyme concentration increased from 0 µg/mL to 5.21 µg/mL, suggesting a near-total bactericidal effect. Additionally, a Pearson correlation of r = -0.8768 was observed between the enzyme concentration and OD600 values, suggesting that ArtE2 has concentration-dependent inhibitory activity. Time-dependent killing curves were generated to study the effect of ArtE2 on E. coli. As illustrated in Figure 5D, the OD600 values began to decrease significantly within 10-15 minutes, and by 20 min post-treatment, the OD600 values had decreased to low levels. Visual inspection of the culture solutions also supports the idea of bacterial cell disruption.

## DISCUSSION

The increasing levels of antibiotic-resistant pathogenic bacteria have become a global public health issue (49). There are estimates of billions of deaths due to these bacteria over the next few decades (50). Many approaches have been investigated as potential solutions to minimize or eliminate the effects of these bacteria on public health (51).

Phages, referred to as endolysins, contain enzymes that have lethal effects on bacteria. Endolysins are hydrolytic enzymes produced by bacteriophages that lyse bacterial cells by degrading peptidoglycans, thereby allowing phage progeny to be released from the cells (52). These enzymes are promising alternative therapeutic agents against antibiotic-resistant bacteria because of their high specificity, rapid action, and low risk of developing resistance (26, 52, 53). We used deep sequencing of *Bacillus altitudinis* and *Bacillus aryabhattai* genomes to identify and isolate a group of genes encoding endolysins. We characterized the isolated endolysins using *in silico* and biological analyses.

All the endolysins analyzed shared the same EADs (N-acetylmuramoyl-L-alanine amidase), indicating that they cleaved the same bond within the peptidoglycan structure (Figure 1A). This finding suggests a conserved mechanism for cell wall degradation by these endolysins. The diversity of CBDs (unknown, SPOR, LysM, peptidoglycan binding, and PlyG) implies that they may attach to bacteria in various ways, exhibit different attachment strengths, and potentially target different bacterial growth stages (e.g., dividing cells for the SPOR domain) (Figure 1A). Endolysins have catalytic domains that target specific bonds in bacterial peptidoglycan (54, 55). Differences in these domains affect substrate cleavage efficiency (56). The targeted endolysin for a particular bacterial cell wall has an optimal configuration, which results in enhanced activity. Another reason for increased enzymatic activity is the presence of cell wall-binding domains (CBDs) located at the ends of each endolysin, these CBDs can bind to receptors on the surface of bacteria (58, 59). As a result of stronger or more specific binding between endolysins and bacterial receptors, enzymatic activity is enhanced, as it maintains longer contact with its target, thereby increasing lysis efficiency (41, 59, 60).

There is considerable variation in the number of bacterial species or strains targeted by some endolysins compared to the number of bacterial species or strains targeted by others (37, 61, 62). Although narrow-specificity endolysins may demonstrate higher activity against a single pathogen, they are less versatile than broad-specificity endolysins (37, 57). In addition to differences in specificity, there are structural differences with respect to the conformation of endolysins, which can explain the varying exposure of either the active catalytic site or binding domain(s) that contribute to both biological activity and specificity (63). Several variations were observed in the endolysins examined in this study. Only seven amino acids were highly conserved in all the endolysin amino acid sequences. Most of these conserved amino acids were localized in the EADs (Figure 1B).

The diversity of CBDs suggests that these endolysins could be tailored for specific applications and that the SPOR domain might be effective against actively dividing Bacillus populations. Endolysins with LysM and/or peptidoglycan-binding domains are highly specific for peptidoglycans. An unknown functional domain may have a less efficient or less specific binding mechanism until its CBD is characterized. The SPOR domain in endolysins plays a critical role in targeting and binding the enzyme to the bacterial cell wall, particularly in gram-positive bacteria (64–68). This binding enhances lytic activity by anchoring it to the substrate, ensuring the efficient degradation of the cell wall, which leads to bacterial lysis (66, 68).

It is likely that these conserved regions of endolysin enzymes are composed of critical functional areas, including the catalytic region, where endolysins break down peptidoglycans in bacterial cell walls. The variable regions may contain host-specific features through the cell wall-binding domain, which determines the specificity of enzyme action. Sequence comparison of endolysins allows inference of evolutionary relationships. When there are very few gaps and an overall high degree of similarity, it is indicative of a recent common ancestry. When there are many gaps and little similarity, it is indicative of a more distant evolutionary relationship owing to the influence of selective forces exerted by the environment on endolysins. The variation in the sequence data of the endolysins clearly indicated that each endolysin targets different bacterial strains within a species.

Clustering may be a useful method for identifying endolysins with broad or narrow activity against selected pathogens and therefore provides information on the evolutionary history of endolysins. Clusters suggested that endolysins have evolved to target different components of bacterial cell walls, which generally differ among Gram-positive, Gram-negative, and mycobacterial strains (Figure 1C). The separation of clusters into distinct groups may indicate a difference in function, possibly due to variations in either the catalytic domain of endolysin, which determines its enzymatic properties, or the cell wall-binding domain, which determines its substrate specificity.

Endolysin phylogenetics demonstrated a complex evolutionary history that contributed to the assortment of endolysins with multiple discrete groups among the 48 endolysins examined in this study, representing a variety of bacterial genera such as *Streptococcus*, *Staphylococcus*, *Mycobacterium*, *Enterococcu*s, *Salmonella*, *Acinetobacter*, *Pseudomonas*, Klebsiella, and *Neisseria* (Figure 1C). The diversity observed in this study provides evidence that several divergent evolutionary pathways have been created in response to different host bacteria. Endolysins derived from *B. altitudinis* and *B. aryabhattai* form a subgroup that is closely associated with endolysins from *Streptococcus*, *Staphylococcus*, and *Mycobacterium*. This clustering of genes may be due to similar structural or functional attributes resulting from either convergent evolution or horizontal gene transfer because the phylogenetic separation between these genera is large.

This finding indicates that endolysins-3 and -7 and *Streptococcus pneumoniae* are linked through a common evolutionary path, possibly via the same binding/lysis mechanism used on different parts of the cell wall of *Streptococcus* (Figure 1C). The variety and specificity of endolysins show great potential for use as targeted antibacterial agents against multidrug-resistant species, such as *Staphylococcus* and *Streptococcus*. However, the effectiveness of these endolysins depends on the range of hosts, as described in the phylogenetic groupings. Interestingly, other endolysins from gram-negative bacteria (*Salmonella*, *Acinetobacter*, *Pseudomonas*, *Klebsiella*, and *Neisseria*) were observed in the main group, suggesting the presence of *B. altitudinis* and *B. aryabhattai* (Figure 1C). Phylogenetic analysis of endolysins was performed using 438 phage genomes and 454 endolysin genes. Among the endolysin sequences, eight different catalytic domains and seven cell-wall-binding domains have been identified (41).

The recombinant expression of *B. altitudinis* and *B. aryabhattai* endolysins allowed the evaluation of their biological activity against the gram-positive bacterium *Staphylococcus aureus*. Endolysin-2, which contains a SPOR domain, exhibited the highest activity (Figure 1D). The SPOR domain is highly specific for peptidoglycan motifs, such as those lacking stem peptides, and helps localize endolysins to vulnerable areas, boosting their antibacterial effectiveness (64–68). It specifically recognizes and binds to peptidoglycan, often at sites of cell wall remodeling or division, such as the septum, where peptidoglycan is newly synthesized or degraded (66, 68). This could explain why endolysin-2 had higher activity than the other endolysins.

In contrast, engineered endolysins modified through directed evolution or fusion with other proteins (outer membrane permeabilizers) can outperform natural endolysins by enhancing their penetration or activity (33–35). This superior activity often stems from a tailored structure, strong target binding, environmental compatibility, and engineered enhancements, making it more effective for specific applications or bacterial targets (18, 33). We created ArtE2 by fusing endolysin-2 with a polycationic peptide (ArtE2) to enhance its ability to penetrate the outer membrane of gram-negative bacteria and to improve its activity against gram-positive bacteria. This modification overcomes the natural barrier posed by the lipopolysaccharide layer in gram-negative bacteria, which typically prevents endolysins from reaching the peptidoglycan layer (18, 33).

Characterization of protein structure is an important step in understanding its function (69). Analysis of the structure of ArtE2 endolysin allowed us to define the different conformations and structures that play an important role in its interaction with the peptidoglycan of the bacterial cell wall (Figure 2A). In contrast, knowledge of the different amino acids conserved within the different classes of endolysins and those that are exposed is important. This allowed us to determine the effectiveness of the interactions with peptidoglycans. The exposed amino acids were highly conserved in the catalytic region of endolysin (Figure 2B).

The sequences contained a mixture of polar (S, T, N, and Q), nonpolar (V, L, I, and A), charged (D, E, K, and R), and aromatic (F, Y, and W) amino acids. Most sequences have a conserved ‘D’ (aspartic acid, negatively charged), which might be part of a catalytic site, as aspartic acid often plays a role in enzymatic activities. There is a conserved ‘G’ (glycine) in most sequences, which is a small, flexible amino acid often found in turns or loops of proteins. Gaps can indicate structural differences or evolutionary divergence among the endolysins (Figure 2B).

Pocket 1 contained a combination of positively and negatively charged residues (charged: ARG; polar: ASP, GLU, SER, THR, and HIS), along with several residues classified as either charged or polar (non-polar: ALA, VAL, LEU, ILE, and PHE). The presence of all three types of residues in this pocket is consistent with the formation of multiple types of interactions with a potential ligand (electrostatic interactions, hydrophobic interactions, and hydrogen bonds). Histidine residues (HIS-10, HIS-79, and HIS-128) have previously been shown to be involved in either the catalytic process or binding of a ligand to an enzyme, as they function both as proton donors and acceptors. Charged residues in Pocket 1 include (in addition to histidines) six negatively charged amino acids (ASP-7, ASP-14, GLU-24, GLU-89, and GLU-141) and one positively charged residue (ARG-63); these residues may participate in salt bridge or hydrogen bonding interactions with a ligand, thereby helping to stabilize it within the pocket. The nonpolar residues found in Pocket 1 (ALA, VAL, LEU, ILE, and PHE) contribute to hydrophobic interactions, which are frequently important for the binding of a ligand to a pocket. In endolysins, residues such as histidine, aspartic acid, and glutamic acid are often part of the catalytic site because they participate in the hydrolysis of peptidoglycans in bacterial cell walls (Figure 3A). This pocket likely corresponds to the active site where the enzyme cleaves the bacterial cell wall (by hydrolyzing peptidoglycan bonds). This may be a minor allosteric site or a non-specific binding region. The residues in pocket 1, particularly catalytic residues such as histidine and aspartic/glutamic acid, support the idea that this pocket is the active site of endolysin. Endolysins typically have a catalytic domain that targets specific bonds in the bacterial cell wall; pocket 1 fits this role (Figure 3A).

The coordinates of pocket 1 were used for molecular docking simulations to predict the binding of substrates or inhibitors to endolysins. The analysis suggested that Pocket 1 was the primary binding site of the endolysin, likely its active site, given its high score, probability, and the presence of catalytically relevant residues, such as histidine, aspartic acid, and glutamic acid. Pocket 2 appeared to be a secondary site, with a lower likelihood of being functionally significant (Figure 3A).

Docking studies have predicted the interactions between ArtE2 and peptidoglycan, which are crucial for understanding its antibiotic function. Specific residues involved in binding and catalysis have been identified, indicating that significant interactions are necessary for the endolysin activity. ArtE2 effectiveness is enhanced by peptidoglycan accessibility in gram-positive bacteria. The model revealed key residues for binding, suggesting potential for engineering improved endolysins. ArtE2 has a high-affinity binding site, which facilitates peptidoglycan recognition and cleavage.

Muramyl dipeptide (MDP) is a component of the bacterial cell wall (70). It is a small peptidoglycan that forms the rigid structure of most bacterial cell walls (70). MDP consist of a sugar molecule (N-acetylmuramic acid) linked to a short peptide. It is found in both Gram-positive and Gram-negative bacteria (71). Histidine in endolysins interacts with MDP through molecular interactions primarily driven by the imidazole side chain. These interactions depend on the protonation state of histidine, which is influenced by local pH and the chemical environment (72). The imidazole ring of histidine can act as a hydrogen bond donor or acceptor, stabilizing the binding of MDP to the active site of the endolysin. At physiological pH, the imidazole ring of histidine is partially protonated, which results in a positive charge. This protonated form can engage in electrostatic interactions with the negatively charged groups of the MDP, enhancing substrate recognition. In its neutral form, histidine can act as a π-system, potentially interacting with the positively charged groups. In some endolysins, histidine may contribute to catalysis rather than to direct substrate binding (Figure 3B, ii).

The distances suggest typical hydrogen bonding (2.5-3.2 angstroms), indicating that certain residues stabilize peptidoglycan in the active site. Histidine (HIS) residues (His-10, His-79, and His-128) may aid the catalytic activity of ArtE2 by cleaving the glycosidic bond. Serine (Ser-77) can form hydrogen bonds and act as a nucleophile. The hydrophobic residues isoleucine (Ile-80) and valine (Val-129) contribute to the binding pocket structure and substrate stabilization. Arginine (Arg-63) may interact ionically with negatively charged groups on peptidoglycan, indicating a polar and potentially catalytically binding site (Figure 3C, ii). These residues play a role in stabilizing the negatively charged peptidoglycan groups.

Endolysins such as ArtE2 typically cleave specific bonds in peptidoglycan monomers, such as the β-1,4-glycosidic bond between N-acetylmuramic acid (NAM) and N-acetylglucosamine (NAG) or peptide cross-links. Histidine residues (His-10 and His-79) may act as catalytic dyads, with one histidine acting as a base to deprotonate a water molecule and the other stabilizing the transition state (Figure 3C ii). The positively charged residues (Arg-63) likely anchor peptidoglycan by interacting with its negatively charged groups, ensuring a proper orientation for catalysis.

Moreover, valine, a non-polar amino acid with a branched isopropyl side chain, interacts with MDP in endolysins through hydrophobic and van der Waals interactions (73). Their nonpolar nature drives the interactions because they lack polar or charged groups. The bulky side chain of valine can form weak van der Waals interactions with MDP atoms, contributing to the binding affinity. The compact branched structure of valine may also play a steric role in shaping the active-site geometry for catalysis. Their nonpolar nature prevents them from forming hydrogen bonds, electrostatic interactions, or catalytic roles (Figure 3B ii). The interaction between a serine residue and the MDP structure involves hydrogen bonding or nucleophilic interactions depending on the endolysin catalytic mechanism. The hydroxyl group of serine can stabilize MDP substrates, and in some endolysins, it may undergo nucleophilic attack, facilitating cleavage (Figure 3B, ii). In the case of isoleucine, endolysin primarily interacts with MDP through hydrophobic or van der Waals interactions. It stabilizes the substrate in the active site and aids in substrate positioning or binding pocket formation. Isoleucine bulky side chains provide weak interactions, supporting active site structural integrity or substrate specificity but are not directly catalytic (Figure 3B, ii).

In contrast, endolysins interact with peptidoglycan monomers, which are repeating disaccharide units cross-linked by peptide chains (74). They target peptide bridges and glycosidic bonds in peptidoglycans, with a catalytic domain that recognizes and cleaves specific bonds (74). Endopeptidases target peptide cross-bridges, breaking the bonds between amino acids such as D-alanine and diaminopimelic acid (75). Endolysins primarily act on polymeric peptidoglycans in cell walls; however, their interaction with isolated peptidoglycan monomers is less efficient (56). *In vitro* studies have shown that endolysins can cleave monomeric units with lower affinity; however, this disrupts the peptidoglycan layer, leading to cell wall weakening and bacterial lysis, particularly in gram-positive bacteria (62).

The interaction between the histidine residue in endolysin and the peptidoglycan monomer structure involves hydrogen bonding, electrostatic interactions, and catalysis (76, 77). It can act as a base, acid, or nucleophile, with a pKa near the physiological pH, allowing it to be protonated or deprotonated (77). The imidazole ring nitrogen atoms can form hydrogen bonds with the polar groups in peptidoglycan, thus stabilizing the substrate binding (78). If protonated, histidine can form ionic interactions with negatively charged groups, enhancing substrate affinity. In many endolysins, histidine acts as a general acid/base in the active site, either deprotonating a nucleophile to attack the glycosidic bond or protonating the leaving group to stabilize the transition state during bond cleavage (Figure 3C (ii)). Interestingly, the interaction between an arginine residue and the peptidoglycan monomer structure involves electrostatic interactions, hydrogen bonding, and substrate stabilization. The positively charged guanidinium group of arginine is highly polar and versatile, making it well suited for interactions with polar and anionic peptidoglycan groups (79, 80). Positively charged arginine can form strong ionic bonds with many negatively charged components in the peptidoglycan layer, including carboxyl groups on the peptide stem and a phosphate-like negative charge on the lactyl group of N-Acetylmuramic Acid (NAM). It is also a multi-hydrogen donor, capable of forming multiple hydrogen bonds with each of the carbonyl oxygen atoms of NAM, as well as with the amide bonds of the peptide backbone and the hydroxyl groups of both NAM and N-acetylglucosamine (NAG) (Fig. 3C, ii).

The recombinant endolysin, ArtE2, exhibited strong and rapid lytic activity against *S. aureus* and *E. coli*. Interestingly, a concentration of 5.21 µg/ml completely inhibited both the bacterial strains at different time points (Figure 5). Interestingly, the same concentration inhibited *E. coli growth* more than *S. aureus* growth. The endolysin ArtE2 with a polycationic peptide showed greater activity against gram-negative bacteria than gram-positive bacteria, primarily due to differences in the bacterial cell wall structure and the interaction of the polycationic peptide with these structures (Figure 5).

Gram-negative bacteria contain a thinner peptidoglycan layer than gram-positive bacteria do. Gram-negative bacteria are surrounded by an anionic outer membrane, which allows polycationic peptides to adhere to and disrupt the bacterial outer membrane (17, 81, 82). Gram-positive bacteria have a thicker peptidoglycan layer, but lack an outer membrane. Therefore, they have a much weaker electrostatic attraction to polycationic peptides than gram-negative bacteria (17, 81–82). The anionic outer membrane of gram-negative bacteria serves as a barrier that limits the ability of endolysins to reach the peptidoglycan layer (81). The outer membrane is disrupted by polycationic peptides, making it easier for endolysins to break down the peptidoglycan layer (17, 83). Although peptidoglycan is directly accessible to Gram-positive bacteria, it reduces the involvement of polycationic peptides in membrane disruption. The positive charge of the polycationic peptide, which originates from arginine and/or lysine residues, can effectively disrupt the outer membrane stability of Gram-negative bacteria, create pores, and increase the permeability of the bacterial outer membrane. This has a much smaller effect on gram-positive bacteria because their cell walls are relatively porous (84).

Studies involving engineered endolysins, such as artilysins, have shown that polycationic peptides significantly increase activity against gram-negative pathogens, such as *Pseudomonas aeruginosa* and *Escherichia coli,* by overcoming the outer membrane barrier (83). Against Gram-positive bacteria, such as *Staphylococcus aureus*, the peptide contribution is less pronounced, as endolysins naturally access peptidoglycan (85). In summary, the polycationic peptide enhanced the endolysin activity against gram-negative bacteria by targeting and disrupting the negatively charged outer membrane, which is a critical barrier. In Gram-positive bacteria, the absence of this barrier reduces the impact of the peptide, as endolysins can act effectively without further membrane disruption.

A specific “Artilysin” combining endolysin KZ144 with the SMAP-29 peptide was highly effective against *A. baumannii* and *P. aeruginosa*, with no observed resistance (86). In addition, a chimeric protein targeting gram-positive streptococci showed enhanced bactericidal activity and stability under varying pH and salt conditions compared with its parental endolysin (18). Engineered from a recombinant library, these artilysins exhibit rapid antibacterial and antibiofilm activity against *P. aeruginosa* (87). Artilysins have shown promise in treating conditions, such as atopic dermatitis, acne, and infected wounds, with case studies reporting significant symptom improvement (26, 88).

Overall, this study demonstrated that the recombinant endolysin ArtE2 is effective for the treatment of infections caused by both *Staphylococcus aureus* and *Escherichia coli* and has good stability. However, to fully assess their potential, additional studies are required to examine their stability, specificity, and effectiveness under various *in vivo* conditions. Endolysins like ArtE2 may offer an attractive alternative to traditional antibiotic treatments; however, there are many factors that have been identified that need to be evaluated, including, but not limited to, delivery systems and stability, host range specificity, immune responses, potential for resistance development, pharmacokinetics, dosing regimens, cost of production, and regulatory issues.

## AUTHOR CONTRIBUTIONS

Orlando Borrás-Hidalgo and Roxana Portieles designed the study, performed bioinformatics analysis, and wrote the manuscript. Roxana Portieles, Xinmin Ma, Jianjian Hu, Hongli Xu, Xiangyou Gao, Nayanci Portal González, Ramon Santos-Bermúdez, Rabia Durrani and Orlando Borrás-Hidalgo performed the experiments. Xinmin Ma and Jianjian Hu assisted in endolysin purification. Orlando Borrás-Hidalgo and Roxana Portieles assisted with data analysis.

## Supplemental Material

**Supplemental Figure S1.** Schematic representation of isolation, identification, recombinant expression, and biological analysis of endolysins.

**Supplemental Figure S2.** Schematic representation of the recombinant expression of endolysin ArtE2.

**Supplemental Figure S3.** Schematic representation of the mechanism of action of the identified endolysins.

**Dataset D1.** List and position of endolysins in *Bacillus altitudinis* and *Bacillus aryabhattai* bacteriophage genomes.

**Dataset D2.** DNA and amino acid sequences of endolysins.

**Dataset D3.** *In silico* analysis of endolysin amino acid sequences.

